# MethylScore, a pipeline for accurate and context-aware identification of differentially methylated regions from population-scale plant WGBS data

**DOI:** 10.1101/2022.01.06.475031

**Authors:** Patrick Hüther, Jörg Hagmann, Adam Nunn, Ioanna Kakoulidou, Rahul Pisupati, David Langenberger, Detlef Weigel, Frank Johannes, Sebastian J. Schultheiss, Claude Becker

## Abstract

Whole-genome bisulfite sequencing (WGBS) is the standard method for profiling DNA methylation at single-nucleotide resolution. Many WGBS-based studies aim to identify biologically relevant loci that display differential methylation between genotypes, treatment groups, tissues, or developmental stages. Over the years, different tools have been developed to extract differentially methylated regions (DMRs) from whole-genome data. Often, such tools are built upon assumptions from mammalian data and do not consider the substantially more complex and variable nature of plant DNA methylation. Here, we present MethylScore, a pipeline to analyze WGBS data and to account for plant-specific DNA methylation properties. MethylScore processes data from genomic alignments to DMR output and is designed to be usable by novice and expert users alike. It uses an unsupervised machine learning approach to segment the genome by classification into states of high and low methylation, substantially reducing the number of necessary statistical tests while increasing the signal-to-noise ratio and the statistical power. We show how MethylScore can identify DMRs from hundreds of samples and how its data-driven approach can stratify associated samples without prior information. We identify DMRs in the *A. thaliana* 1001 Genomes dataset to unveil known and unknown genotype-epigenotype associations. MethylScore is an accessible pipeline for plant WGBS data, with unprecedented features for DMR calling in small- and large-scale datasets; it is built as a Nextflow pipeline and its source code is available at https://github.com/Computomics/MethylScore.

## Introduction

Cytosine methylation, which is often used synonymously with DNA methylation, describes the covalent attachment of a methyl group to carbon 5 of cytosine, resulting in 5-methylcytosine (5mC). In plants, as in most eukaryotes, DNA methylation is part of an epigenetic mechanism involved in transposon silencing (Miura et al., 2001), heterochromatin formation (Lippman et al., 2004), and gene regulation (Jaenisch and Bird, 2003). It also plays a role in genome organization (Zemach et al., 2013; Huff and Zilberman, 2014), regulation of development (Finnegan et al., 1996; Papareddy et al., 2021; Ronemus et al., 1996) and imprinting (Jullien et al., 2008; Pignatta et al., 2018; Gehring et al., 2006). DNA methylation can also be a source of phenotypic variation in natural populations (Schmitz et al., 2013; Eichten et al., 2011; Heyn et al., 2013).

Despite recent advances in long-read sequencing, short-read WGBS is still considered the gold standard for analysis of DNA methylation at single-base resolution. Treatment of genomic DNA with sodium bisulfite causes hydrolytic deamination of unmethylated cytosine (C) into uracil, which upon sequencing library amplification by PCR is converted to thymine (T) (Frommer et al., 1992). The initial step is kinetically inhibited in the case of 5mC, and so methylated cytosines will be interpreted normally during short-read sequencing, whereas unmethylated, converted cytosines will be interpreted as T by the base caller. Comparison to the known reference genome can subsequently reveal the methylation state of each cytosine.

### Features of plant DNA methylation

In many eukaryotic organisms, cytosines can be in a methylated state when they are in a CG sequence context, i.e., followed by a guanine (G). In mammalian genomes, for example, the majority of CGs are methylated, while unmethylated cytosines are grouped in so-called CpG islands, which are non-randomly distributed along the genome and play an important role in transcriptional regulation (Deaton and Bird, 2011). In plant genomes, however, the situation is fundamentally different and can vary substantially from one species to another. DNA methylation in plants occurs in three possible sequence contexts: CG, CHG, and CHH (where H is any base but G) (Law and Jacobsen, 2010). Each type of methylation is established, maintained and regulated by a specific molecular machinery (Stroud et al., 2013b; Henderson and Jacobsen, 2007; Law and Jacobsen, 2010). The frequency of methylation along the genome differs by sequence context; for example, in the model plant *Arabidopsis thaliana*, CGs are methylated most frequently relative to the total number of cytosines in that context (∼24% of all CGs are 5mCGs), followed by CHG (∼7%) and CHH (∼1.7%) (Cokus et al., 2008).

While DNA methylation in CG and CHG is symmetric, i.e., the complementary sequences on the opposite strand mirror the methylation state, CHH methylation is asymmetric and occurs on just one strand at that particular locus. For CG methylation, the information retained by semi-conservative DNA replication facilitates re-establishment of methylation on the daughter strand via the methyltransferase METHYLTRANSFERASE 1 (MET1) that, akin to its mammalian homologue DNA METHYLTRANSFERASE 1 (DNMT1), shows substrate preference to hemimethylated DNA (Kankel et al., 2003; Jeltsch, 2006; Saze et al., 2003). In contrast, maintenance of CHG and asymmetric CHH methylation depends on additional factors such as histone modifications and small RNAs. Methylation of CHG cytosines by CHROMOMETHYLASE 3 (CMT3) reciprocally depends on H3K9 dimethylation (Jackson et al., 2002; Du et al., 2012), while CHH methylation requires CHROMOMETHYLASE 2 (CMT2) (Stroud et al., 2014; Zemach et al., 2013) and DOMAINS REARRANGED METHYLTRANSFERASE2 (DRM2) recruited through small-RNA-dependent RNA-directed DNA methylation (RdDM) (Aufsatz et al., 2002; Cao et al., 2003; Cao and Jacobsen, 2002); (Wassenegger et al., 1994; Matzke and Mosher, 2014).

It is important to note that — at least in some plant species, including *A. thaliana* — DNA methylation in the different sequence contexts also differs in terms of methylation rate, i.e., in the consistency of the methylation status of a given cytosine across different cells. CG cytosines tend to have a binary methylation state, being either always unmethylated or almost always methylated in different cells of a given tissue. In contrast, cytosines in CHG and CHH show more variable methylation status, with mean methylation rates across all methylated CHG and CHH cytosines of ∼50% and ∼30%, respectively. This has consequences for the analysis of differential methylation between samples (Becker et al., 2011), for example in software which would model the underlying methylation status based on expected distributions (Hansen et al., 2012; Hebestreit et al., 2013; Korthauer et al., 2018).

### Whole-genome DNA methylation analysis as a means to understand natural variation and stress response

Since the first characterizations of the whole-genome DNA methylation profile in *A. thaliana* (Cokus et al., 2008; Lister et al., 2008), there has been a growing interest in studying this epigenetic mark at the genomic level to better understand developmental processes (Pignatta et al., 2018; Manning et al., 2006), stress responses (Wibowo et al., 2016; Liu et al., 2018), phenotypic plasticity, and natural variation (Schmitz et al., 2013; Kawakatsu et al., 2016). For example, heritable genetic variation alone is not always sufficient to explain the range of phenotypic diversity observed for individuals of the same species, and epigenetic variation is likely to account for at least some of the missing heritability (Manolio et al., 2009). Epigenetic variation, albeit often confounded by genetic variation, has been recognized as a source of natural diversity (Riddle and Richards, 2002; Cervera et al., 2002; Vaughn et al., 2007). As a consequence, some phenotypic traits, including floral symmetry (Cubas et al., 1999), fruit development and morphology (Manning et al., 2006; Ong-Abdullah et al., 2015; Zhong et al., 2013), plant height (Miura et al., 2009), and resistance to pathogens (Liégard et al., 2019), have been associated with naturally occurring epigenetic alleles (epialleles) that do not appear to be due to linked genetic variation. In addition, studying the conditional dynamics of DNA methylation at the whole-genome level has highlighted the contribution of epigenetic regulation to stress response and tolerance (Wibowo et al., 2016).

### The challenges in determining differential DNA methylation

DNA methylation is highly dynamic, and there is substantial variation among individuals or (sub)populations. It is important to distinguish specific, relevant differences from stochastically occuring methylation differences between biological replicates. On a genome-wide scale, this is challenging without *a priori* knowledge of stratification, either based on sequence context or by sample grouping. Most experimental studies contrast DNA methylation from different samples to each other, be it mutant background and wild type, treatment and control, or different natural accessions. Statistical comparisons between samples aim to identify DNA methylation differences at either the single-cytosine (differentially methylated positions; DMPs) or the region (differentially methylated regions; DMRs) level. While DMPs provide useful information on the rate at which epigenetic changes occur (Becker et al., 2011; Schmitz et al., 2011; van der Graaf et al., 2015), DMRs are arguably more relevant in a functional biological context because they can affect contiguous stretches of DNA and hence potentially influence the accessibility of regulatory elements. However, the nature of WGBS data imposes several caveats to accurately determining DMPs and DMRs. Some of these reside in the experimental design or the quality of the sequencing library. Insufficient replication, for example, limits the statistical power of differential analyses. Uneven coverage can bias the base calling because of sequencing error rates, while incomplete bisulfite conversion can cause a false estimate of methylated cytosines, as can duplicated reads that arise from library over-amplification.

The statistical analysis of differential methylation often brings along another set of caveats. For example, strategies that are based on defining DMRs as clusters of spatially adjacent DMPs, such as *DSS* (Feng et al., 2014; Ziller et al., 2013; Schultz et al., 2015; Feng et al., 2014), are subject to a heavy multiple testing burden resulting from the large number of cytosines in the genome that need to be tested individually. The same applies to window- or sliding-window-based approaches, as implemented, for example, in *methylKit* (Akalin et al., 2012). In contrast, strategies that call DMRs only in pre-defined regions, e.g., in annotated features, risk missing relevant loci in the analysis.

More recently developed tools for DMR calling are centered around pre-selection of genomic regions. *metilene* (Jühling et al., 2016), *dmrseq* (Korthauer et al., 2018) and *HOME* (Srivastava et al., 2019) implement multi-step strategies that first restrict the testable genome space to candidate regions with evidence of methylation, prior to assessing significant differences. Even though *HOME* takes non-CG methylation into account, all three of these tools have been developed with mammalian (mostly human) DNA methylation data in mind and are therefore built on assumptions, including most cytosines being methylated, strong local correlation between methylation states, predominant (or exclusive) CG methylation, and largely binary methylation states, that do not reflect the distinct genetic control underlying the CG, CHG and CHH methylation contexts in plants. This implies that if these assumptions are indiscriminately applied, many methylated cytosines and regions are likely to be falsely discarded.

Here, we set out to address this gap by developing a tool for the robust identification of DMRs from plant WGBS data that takes into account the complexity and variability of plant DNA methylation, while using an informed, restricted set of candidate regions for statistical testing. Furthermore, we aimed to avoid the necessity for pre-defining sample groups, required by existing tools, to increase the applicability to sample populations with inherent group structure and to prevent experimenter bias. We present MethylScore, a convenient and stable pipeline that enables biology researchers with little computational background to process WGBS data, from read alignments to DMR output. The differential methylation analysis module of MethylScore is built around the two-state Hidden-Markov-Model-based approach described by (Molaro et al., 2011). To identify and segment methylated regions in plant genomes, independent of prior information, we extended the original implementation beyond the CG sequence context, allowing the algorithm to train distinct parameter sets for each methylation context.

MethylScore trains on the different properties of CG, CHG, and CHH methylation by estimating parameters of a beta-binomial distribution for each sequence context, accounting for both stochastic variance in the coverage distribution (assumed to be beta distributed) as well as between-sample biological variance (binomially distributed). This means that MethylScore learns from the actual WGBS data, thus avoiding potentially erroneous or species-specific assumptions about the distribution of methylation or arbitrary thresholds. This way, the number of statistical tests is constrained to regions of interest, curtailing the multiple-testing problem. Moreover, its built-in population-scale approach allows identifying DMRs in large datasets, without prior information on sample groups, enabling an unbiased DMR calling. Using publicly available datasets from *A. thaliana* (1001 Genomes Consortium, 2016; Kawakatsu et al., 2016; Tedeschi et al., 2019; Ning et al., 2020; Zhang et al., 2021; 1001 Genomes Consortium, 2016; Kawakatsu et al., 2016; Wibowo et al., 2018) and rice (Stroud et al., 2013a), we show that MethylScore is able to segment plant genomes with very different global DNA methylation profiles. In absence of sample information, MethylScore identified group-specific DMRs and was able to detect population signals in datasets with hundreds of samples. Ultimately, we used the DMRs thus identified in the *A. thaliana* 1001 Genomes and Epigenomes datasets (1001 Genomes Consortium, 2016; Kawakatsu et al., 2016) to detect known and unknown genotype-epigenotype associations. MethylScore is built as a Nextflow pipeline, allowing ease-of-access and broad usability; the pipeline can be downloaded from https://github.com/Computomics/MethylScore.

## Results

### Outline of the modular MethylScore pipeline to call DMRs from plant WGBS data

MethylScore is implemented as a modular pipeline that is built in Nextflow (Di Tommaso et al., 2017) for maximal ease-of-access (Figure 1; Materials and methods). It uses reference-aligned sequences in *bam* format or tabular single-cytosine information in *bedGraph* format as they are produced by common WGBS primary analysis tools such as *bismark* (Krueger and Andrews, 2011) or *MethylDackel* (*https://github.com/dpryan79/MethylDackel*). The MethylScore pipeline performs the following general steps: (*i*) calling of methylated cytosines and determining the per-cytosine methylation rate, (*ii*) identifying contiguous regions with high DNA methylation in each sample, (*iii*) determining segments that for statistical testing, and (*iv*) performing statistical tests to identify DMRs. To start the pipeline, users need to provide a two-column sample sheet with sample identifiers and file paths to the respective *bam* or *bedGraph* files as well as to the reference genome sequence in *fasta* format. If multiple lines in the sample sheet share the same sample identifier, MethylScore will assume technical replication and merge the files prior to further processing.

**Figure 1.**
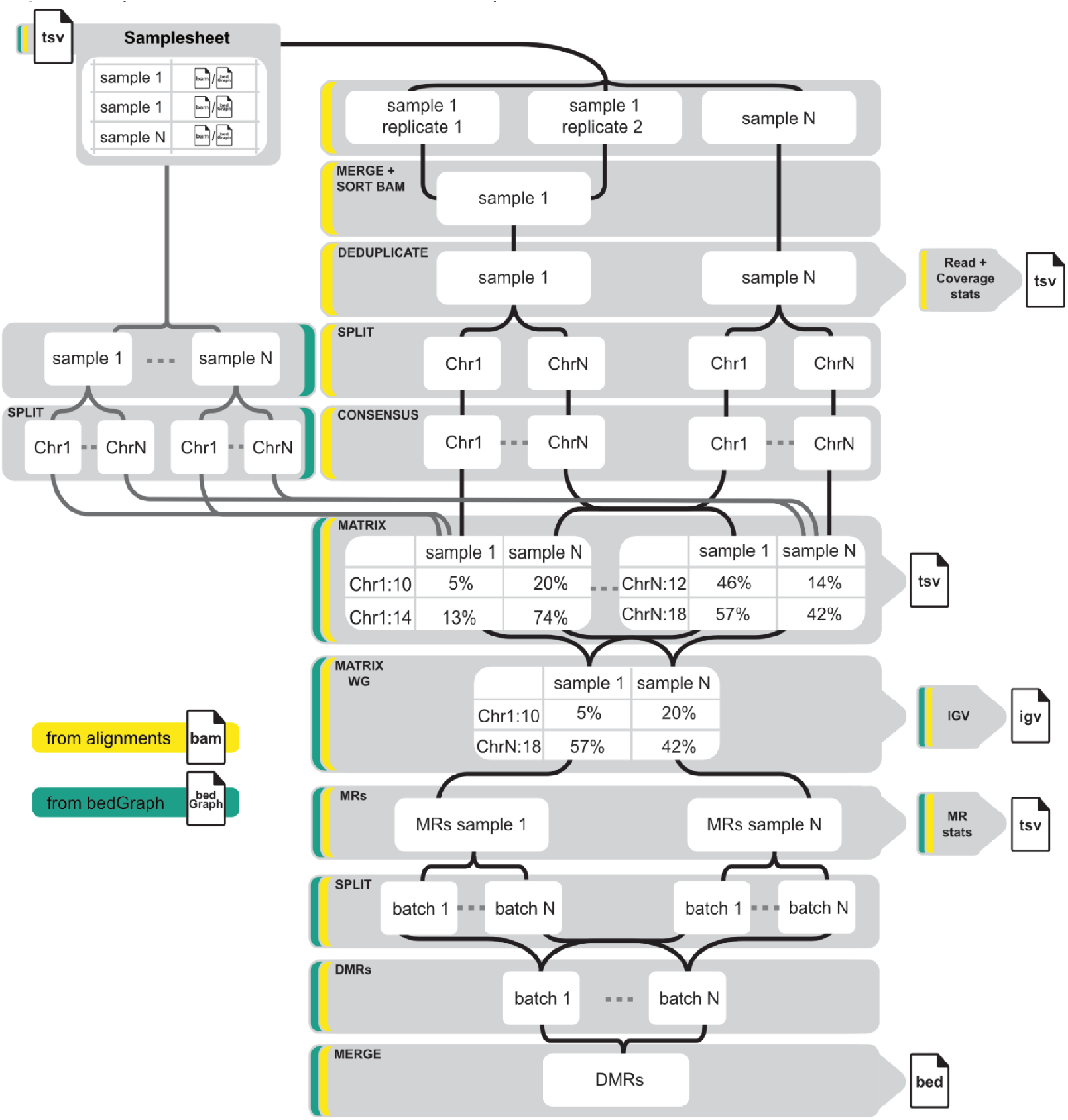
Schematic overview of the MethylScore pipeline. Schematic overview of processing steps (grey). Different workflow entry points based on the type of input data are indicated as coloured borders. The workflow can be started from genomic alignments (*BAM* format; yellow) or tabular single-cytosine methylation information (*bedGraph* format; green). Chr: chromosome; DMR: differentially methylated region; MR: methylated region.

First, the *samtools sort* (Li et al., 2009) function sorts the alignments by genomic coordinates. As many library preparation protocols for bisulfite sequencing incorporate a PCR amplification step during the protocol, alignments can be optionally processed with *picard MarkDuplicates (Picard toolkit, 2019)* to avoid potential bias by non-uniform read amplification. For the next steps up to the methylated region calling, the alignments are split by chromosome or contig, facilitating full parallelisation of downstream processing, which is open to user configuration in order to suit the available computational resources. *MethylDackel extract* (*https://github.com/dpryan79/MethylDackel*) is used for tabulation of single-cytosine information per sample.

The consensus files for each sample and chromosome are merged into a uniform file format containing methylation information for all cytosines across all samples, together with information about sequence context and strand information for each cytosine. After merging individual chromosomes, the resulting genome matrix file is compressed with *bgzip* and indexed with *tabix*, allowing for targeted queries of regions from the compressed file. The genome matrix serves as the input for iterative training of one Hidden Markov Model (HMM) per sample. It is implemented as a two-state (unmethylated/methylated) model, designed to learn three context-specific methylation rate distributions by fitting distinct beta-binomial distributions, thus reflecting the underlying biologically distinct control of these sequence contexts in plants (see Materials and methods).

Based on the resulting parameterization of the trained model, the genome space is segmented into methylated and unmethylated regions by probabilistic classification, yielding genomic coordinates of MRs in *bed* format. These regions are split into batches to facilitate the parallel processing of the ensuing statistical testing strategy. Candidate DMRs in MR batches are preselected based on changes in MR frequency across the sample population for the detection of plausible region boundaries (Supplementary Figure 1). Using an iterative *k*-means algorithm to determine the most suitable number of groups (MacQueen, 1967), samples are clustered according to their mean methylation rates within candidate DMRs. Clusters for each sequence context are assessed for statistical significance by employing log-likelihood ratios based on observed read counts under the beta-binomial assumption and testing against a chi-squared distribution with six degrees of freedom as outlined in (Hagmann et al., 2015), followed by FDR control using Storey’s method (Storey and Tibshirani, 2003). In a last step, DMRs that pass the FDR-corrected significance threshold are merged, yielding one *bed* file per methylation context containing DMRs between sample clusters.

### Segmentation of plant genomes with different DNA methylation composition

We first wanted to explore how MethylScore would segment plant genomes with very different global DNA methylation configurations. In *A. thaliana*, DNA methylation is unevenly distributed, with most methylated cytosines located in the centromeric and pericentromeric regions (Niederhuth et al., 2016). Overall, 11% of cytosines are methylated, with 31.7% of CG, 16.3% of CHG, and 6.2% of CHH cytosines showing methylation rates above zero. In contrast, in the genome of rice (*Oryza sativa*), DNA methylation is more evenly distributed along the chromosomes and more frequent in general, showing 48% CG, 25.3 % CHG and 5.7% CHH methylation. When segmenting WGBS data for both species, MethylScore detected 42,478 methylated regions (MRs) in *A. thaliana* vs. 379,227 in rice, covering 19.5% and 32.5% of the total genome sequence, respectively. Accordingly, the median length of MRs in Arabidopsis (111 bp) was much shorter than in rice (188 bp) (Supplementary Figure 2).

### *MethylScore identifies differentially methylated regions without* a priori *sample information*

We designed MethylScore to be unaware of sample relationships, which means that it does not require any *a priori* information about replicate groups and that the analysis will not be biased by potentially inaccurate assumptions about sample similarities. Instead, for each candidate DMR, MethylScore groups all samples using an iterative *k*-means approach (see Materials and methods), followed by a beta-binomial-based test to statistically assess whether the group means are significantly different from each other. As a result, grouping may not necessarily reflect the preconceived biological design of the experiment for each candidate DMR, yet local grouping of DMRs has the advantage that it allows to identify patterns in the underlying sample structure that might be overlooked when defining groups beforehand. Averaged over all candidate DMRs, methylation differences should be apparent between, for example, treatment groups or developmental stages *if* the underlying hypothesis for these groups to differ in DNA methylation was correct. To test if MethylScore accurately differentiates between-group and within-group differences in this unsupervised approach, we applied it to *A. thaliana* datasets that had previously been described as having substantial levels of between-group methylation divergence.

We first analyzed a relatively simple WGBS dataset of Columbia-0 (Col-0) wild type and two loss-of-function mutants of *EFFECTOR OF TRANSCRIPTION 1* (*et1-1*) and *EFFECTOR OF TRANSCRIPTION 2* (*et2-3*), respectively, as well as of an *et1-1 et2-3* double mutant, with three biological replicates per genotype. These mutants had previously been shown to have local DNA methylation changes compared to wild-type but also compared to each other (Tedeschi et al., 2019). When running MethylScore on all samples without group replicate information, methylation differences in DMRs accurately reflected the genotypic relationship, indicating that the grouping of samples over all DMRs was driven by methylation differences between genotypic groups (Figure 2). In accordance with the original publication, grouping was most conspicuous in CG and CHG contexts (Figure 2).

**Figure 2:**
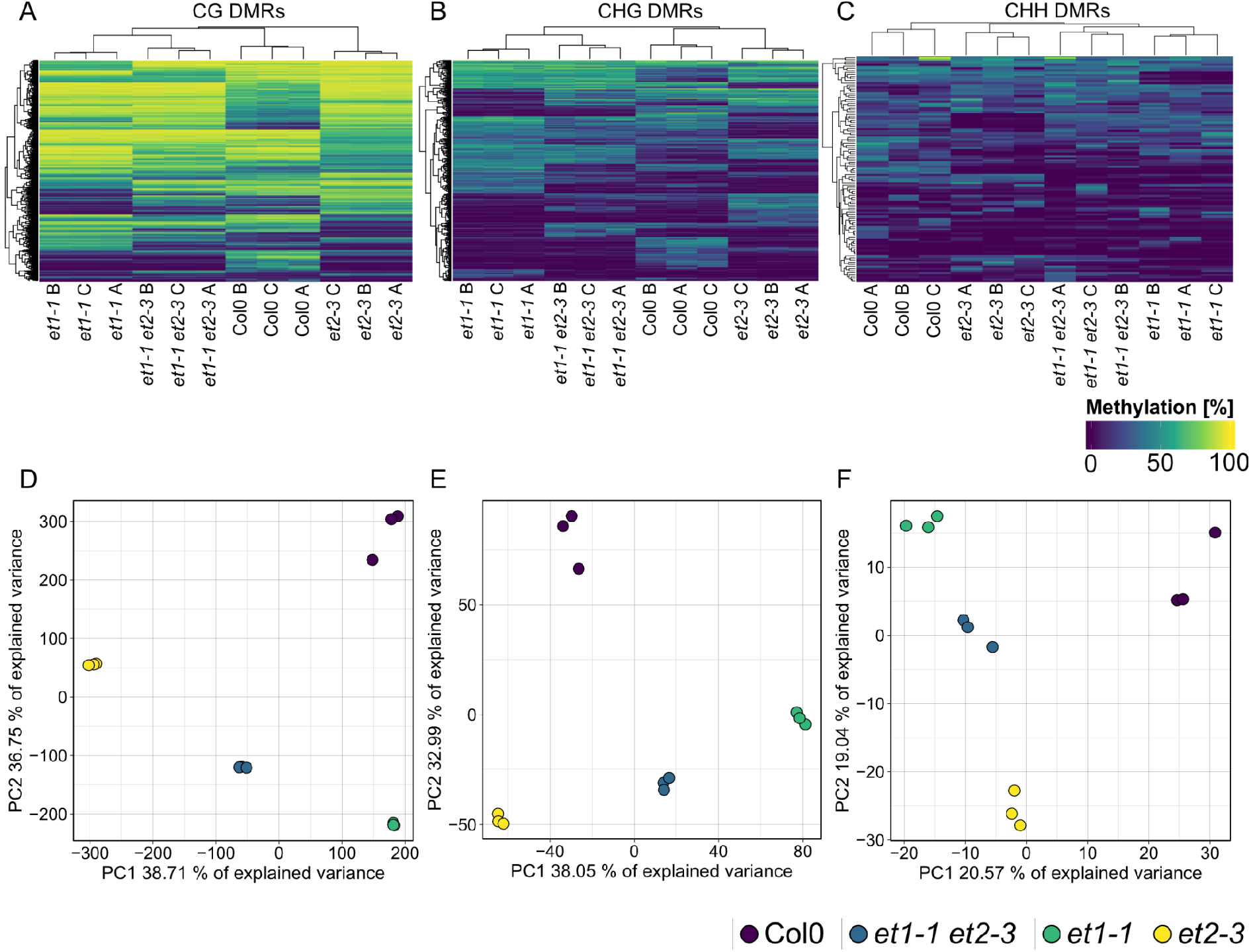
MethylScore DMR calling clusters samples according to genotype. Heatmaps and Principal Component Analyses (PCAs) of mean methylation rates in 9,487 CG- (**A**,**D**), 1,282 CHG- (**B**,**E**) and 741 CHH- (**C**,**F**) context-specific DMRs identified by MethylScore from WGBS data from flowers of *A. thaliana* Col-0 wild-type, respective *et1-1* and *et2-3* single mutants, and *et1-1 et2-3* double mutants. Original data from Tedeschi et al. (2019) (ENA accession PRJEB12413).

Next, we asked whether MethylScore would accurately classify methylation pathway mutants with more pronounced DNA methylation differences to wild-type. Some of these mutants have substantial global loss of cytosine methylation; we therefore wanted to see whether the training of the HMM, which takes into consideration the methylation rate distributions of each sample, might be affected by this. We used a dataset comprising loss-of-functions mutants of the CHG-specific DNA methyltransferase CMT3, the chromatin remodeler DECREASE IN DNA METHYLATION 1 (DDM1), two regulators of these genes, TESMIN/TSO1-LIKE CXC DOMAIN-CONTAINING PROTEIN 5 (TCX5) and TCX6, and of combinations thereof (Ning et al., 2020). Similar to the *et* mutant analysis (Figure 2), MethylScore accurately clustered the samples according to genotype based on DNA methylation rates within DMRs (Figure 3). Context-specific DMRs also resolved CG- and CHG-specific loss of methylation in *ddm1* and *cmt3*, respectively (Figure 3), indicating that deviations from the standard methylation rate distributions did not affect MethylScore analysis.

**Figure 3:**
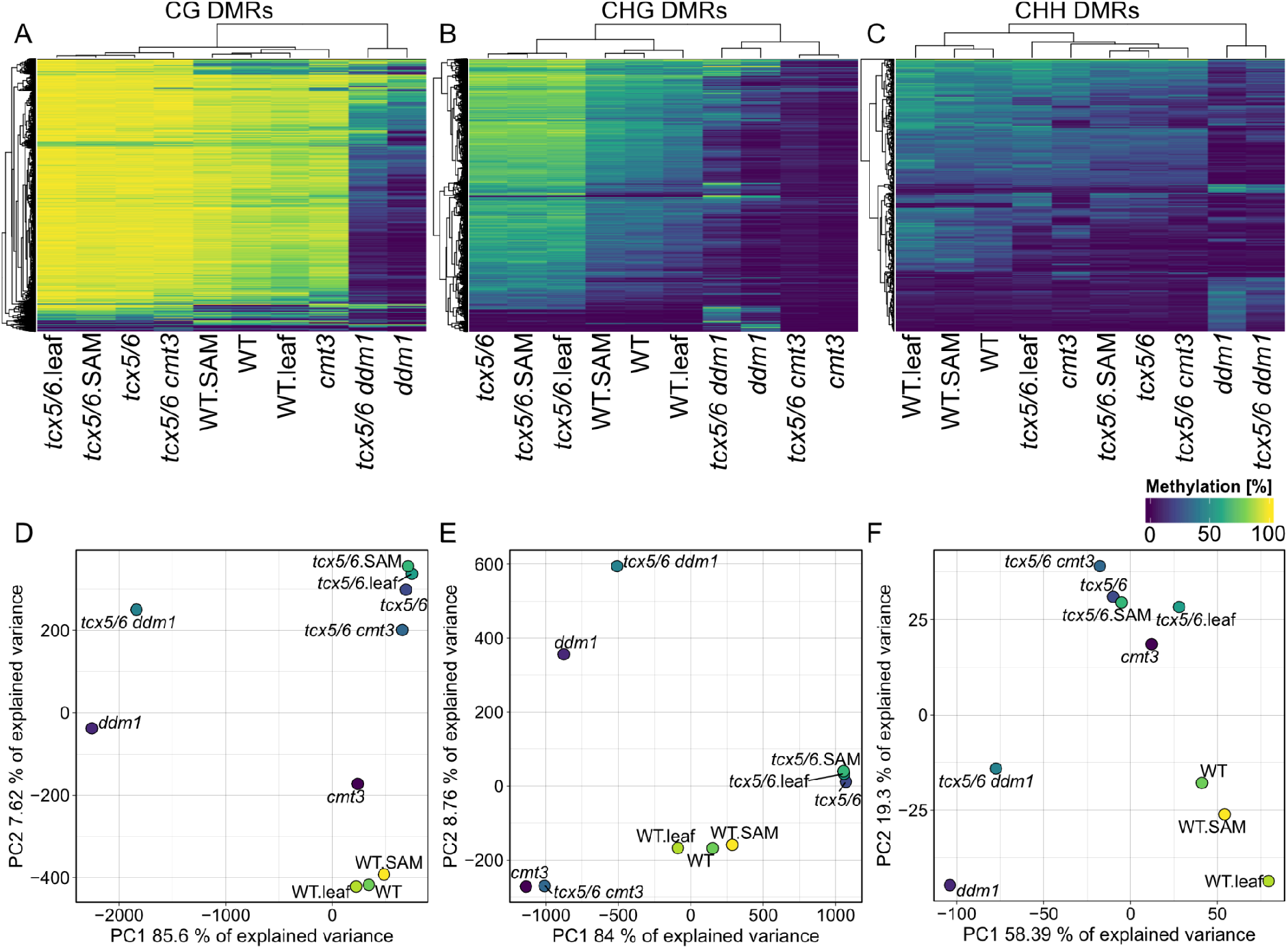
Unsupervised DMR calling from WGBS data of DNA methylation pathway mutants. Heatmaps and PCAs of mean methylation rates in 59,153 CG- (**A**,**D**), 63,385 CHG- (**B**,**E**) and 440 CHH- (**C**,**F**) context-specific DMRs identified by MethylScore from WGBS sequencing data of DNA methylation pathway mutants. The dataset included *ddm1* and *cmt3* single mutants, *tcx5/6* double mutants, as well as *tcx5/6 ddm1* and *tcx5/6 cmt3* triple mutants. DNA had been sampled from leaves and shoot apical meristem (SAM). Original data from Ning et al. (2020); GEO accession GSE137754.

### Clustering of methylated regions across regenerated A. thaliana populations underlines partial maintenance of organ-specific methylation profiles in CG

Mutants known to be affected in DNA methylation arguably are expected to present relatively simple cases for genome-wide differential analyses. To test more complex situations, we applied MethylScore’s population-scale approach to more complex datasets with less predictable group structure. We re-analysed a published dataset of *A. thaliana* regenerants derived from different tissues of origin that had been obtained by induction of somatic embryogenesis (Wibowo et al., 2018). Instead of employing a pairwise testing strategy between condition groups as it had been pursued in the original study, we used MethylScore to preselect candidate regions based on MR frequency changes across the sample population, and hypothesized that such a strategy should naturally find clusters of regenerants with a similar epigenetic setup, thus allowing to differentiate between organ-derived methylation signatures. The data contained three potentially interacting factors that could contribute to epigenetic variation: regeneration via somatic embryogenesis *vs*. sexual reproduction; tissue of origin of the somatic embryos (RO: root origin; LO: leaf origin); and tissue of DNA sampling (leaf or root) in the progeny of the regenerants.

Hierarchical clustering of methylation rates in 1,282 CHG- and 741 CHH-DMRs revealed tissue type during DNA sampling as the major determinant of methylation within these regions (Figure 4A,B). Tissue of origin was subordinated to sampling tissue type. Principal component analysis (PCA) of methylation rates within the same regions also showed clear ordination based on sampling tissue type along PC1 (Figure 4B), for both CHH and CHG contexts. In contrast, among all 9,487 DMRs identified in CG context, tissue of origin appeared to play a role alongside sampling tissue type. Our analysis also recapitulated the main finding of the original publication, namely that DNA methylation in leaves of RO progenitors was more similar to that in root tissue. Our results generally supported the original conclusion of partial maintenance of epigenetic marks retained from the tissue of origin across generations of sexually reproduced offspring, yet suggested CG methylation to be the main driver of this observation (Figure 4A).

**Figure 4:**
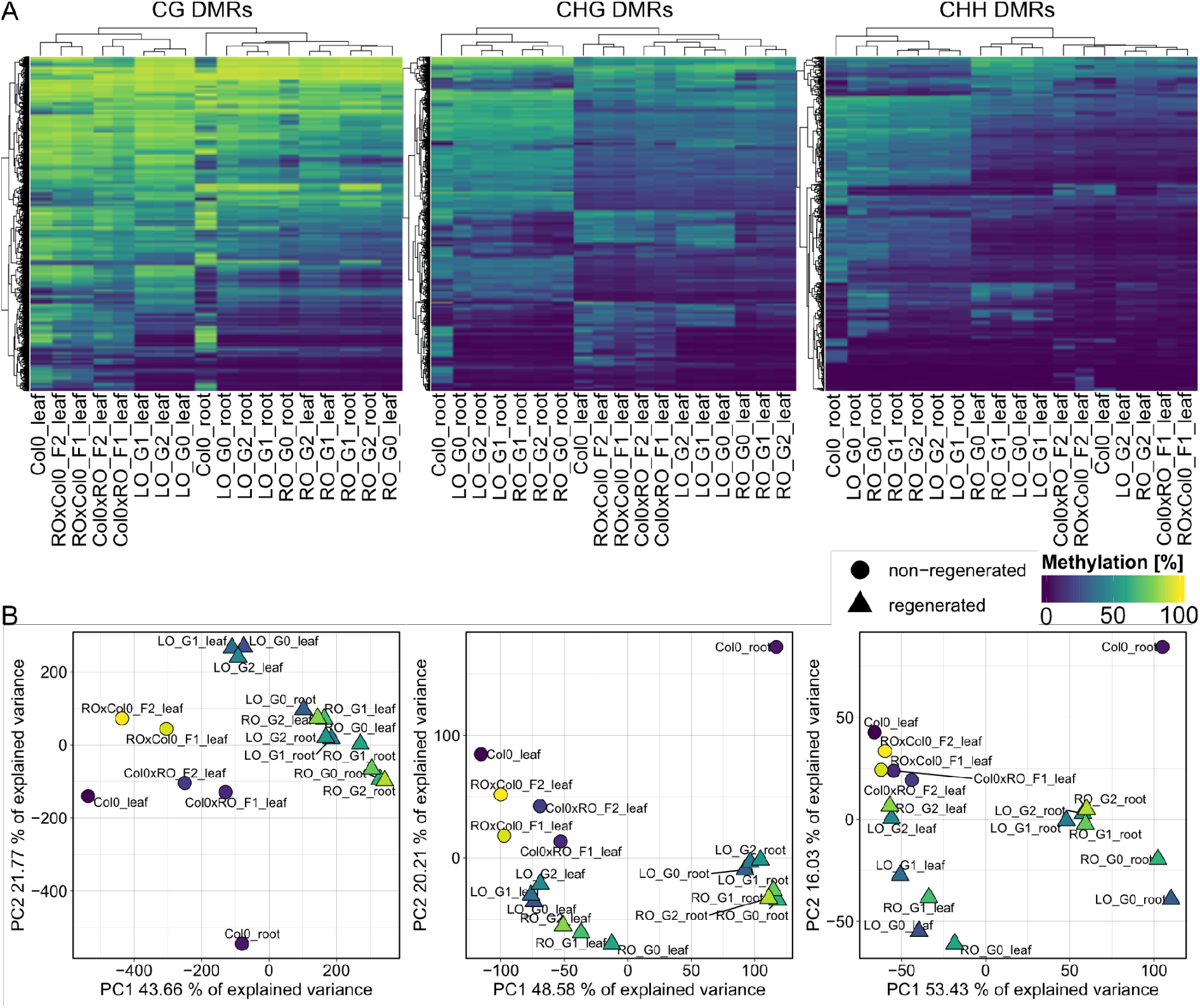
MethylScore population clustering partially reflects epigenetic origin of regenerated plant lineages. **A)** Heatmaps show methylation rate averages in regions identified as differentially methylated in CG, CHG, or CHH methylation contexts (from left to right). The dataset includes leaf and root tissue from Col-0 control plants as well as from generation 1 (G1) and generation 2 (G2) progeny of somatic regenerants from root origin (RO) and leaf origin (LO) somatic embryos, and leaf tissue from F1 and F2 backcrosses of RO and LO regenerants to Col-0. **B)** PCAs for each methylation context using the same data are shown in **A**. Original data from Wibowo et al. (2018), ENA accession PRJEB26932.

To validate whether MethylScore could accurately resolve complex data structures in larger plant genomes with much denser DNA methylation profiles, we also re-analysed a regeneration-related dataset from rice (Stroud et al., 2013a). Similar to the *A. thaliana* dataset, MethylScore identified DMRs that separated regenerated from non-regenerated samples (Supplementary Figure 3). Interestingly, however, clustering of samples based on methylation rates within DMRs changed considerably depending on the sequence context. For example, callus samples were similar to control plants and regenerants in CG-DMRs but stood out as hypermethylated in the CHH context (Supplementary Figure 3).

### DMR calling on a population scale can identify unknown underlying sample structures

MethylScore does not require information on sample grouping but instead clusters samples for each DMR based on methylation rates, so we expected it to be able to call DMRs even in very large population-scale datasets. We also wanted to explore how combined epigenetic and genetic variation would affect the algorithm. Natural genetic and epigenetic variation in *A. thaliana* has been cataloged in the 1001 Genomes and Epigenomes Project (www.1001genomes.org) (1001 Genomes Consortium, 2016; Kawakatsu et al., 2016). We analyzed a subset of 645 *A. thaliana* accessions that had been sequenced by WGBS at the SALK Institute, San Diego, USA. Applying the population-scale approach of MethylScore, we identified 60,797, 16,627, and 8,406 DMRs in CG, CHG, and CHH, respectively. PCA on the accession-specific methylation rates in these regions revealed geographic clustering that separated a cluster of Central Asian accessions from the rest (Figure 5). This grouping occurred for all three sequence contexts and had not been described in the original publication, indicating that MethylScore’s approach can detect data structures that remain hidden with conventional DMR calling tools. Moreover, CG, but not CHG or CHH DMRs, resolved a latitudinal transect for European accessions (Figure 5). We wanted to explore the source of the signal further and used the geographic coordinates of collection sites for each of the *A. thaliana* accessions to retrieve bioclimatic variables from the WorldClimate dataset (www.worldclim.org) (Fick and Hijmans, 2017). While the Central Asian cluster showed strong correlation with low annual average temperature (bio1), strongest correlation was observed for the lowest temperature in the coldest month of the year (bio6). While local climate is strongly confounded by geographic location and thus population structure, this observation is in line with previous reports that showed interdependence between DNA methylation and ambient temperature (Dubin et al., 2015) as well as seasonality (Shen et al., 2014).

**Figure 5:**
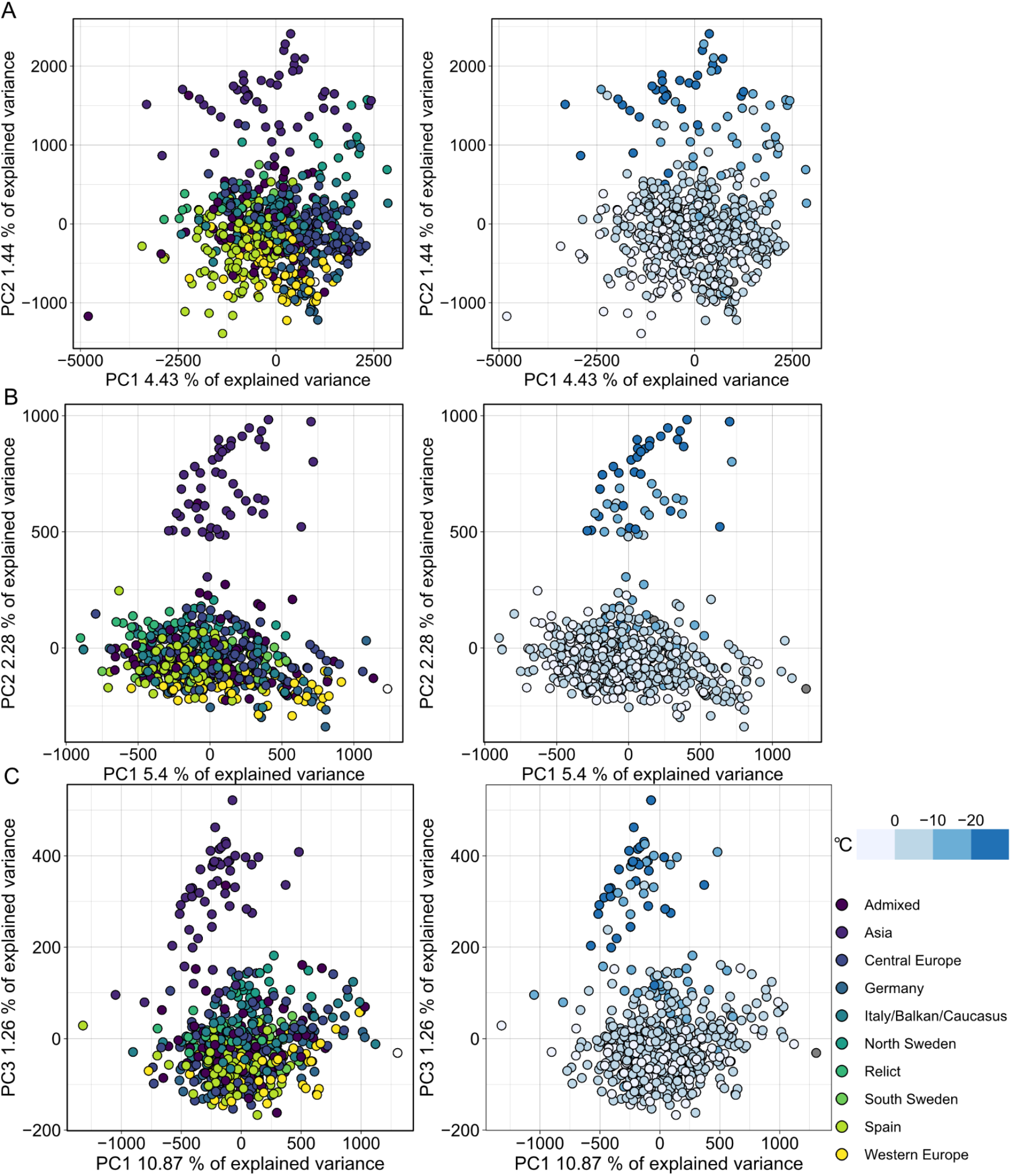
Population structure analysis of natural *A. thaliana* accessions based on DMRs identified by the population-scale clustering approach of MethylScore. PCA shows group formation in CG (A), CHG (B) and CHH (C) methylation contexts. Colors indicate admixture groups (left column) and seasonality with regard to the lowest temperature in the coldest month (right column). Data was retrieved via geographic coordinates of collection sites for each accession from the worldclim.org bio6 dataset (Fick and Hijmans, 2017). Original WGBS published in Kawakatsu et al. (2016).

To further explore MethylScore’s capability to resolve hidden sample/population structures, we applied it to WGBS data of 169 epigenetic recombinant inbred lines (epiRILs) (Zhang et al., 2021). These lines are derived from a cross of Col-0 wild-type and a loss-of-function mutant of DDM1 and propagated by single-seed-descent. Genetically, these lines can be considered near-isogenic; previous studies showed that these epiRILs carry marked differences in DNA methylation in mosaic-like patterns of DNA methylation, some of which are stably inherited through generations of single-seed descent (Johannes et al., 2009; Colomé-Tatché et al., 2012). As every epiRIL has been propagated independently and should therefore be independent from every other line, we did not expect any sample relationships. However, when analyzing the DMRs returned by MethylScore’s population-scale analysis in a PCA, the epiRILs formed several clusters (Supplementary Figure 4). In all three sequence contexts, most lines were contained in one large cluster, with the exception of a few lines that clustered separately (Supplementary Figure 4). This clustering may reflect the fact that lines were selected for WGBS based on phenotypic information (i.e., selected epigenotyping) or yet unknown sources of sample stratification, including genetic structure due to shared TE insertion profiles (Quadrana et al., 2019) in a subset of lines.

### *MethylScore DMRs reveal recurrent trans-acting association signals in* A. thaliana *natural accessions*

MethylScore’s *k*-means clustering leads to grouping of samples into only a small number of groups (mostly two or three), hence within-group variance becomes relatively large when sample numbers are high, as was the case for the 1001 Genomes data. As a result, DMRs with noticeable between-group differences become rare, explaining the relatively low number of DMRs we observed in the 1001 Genomes data, in relation to the size of the dataset. To increase the statistical power to detect genotype-epigenotype associations in a genome-wide association (GWA) mapping approach that uses methylation rates in each DMR as the phenotype vector, we changed from a population-scale to a pairwise DMR calling, comparing each of the 645 accessions individually to the Col-0 reference. On average, each pairwise comparison identified 1,708, 778, and 3,625 DMRs in CG, CHG, and CHH contexts, respectively. Many DMRs from different pairwise comparisons partially or fully overlapped, so we created sets of unions of overlapping DMRs for each context, resulting in 18,044 CG-, 9,674 CHG-, and 25,350 CHH-DMRs.

We queried methylation rates per accession and conducted GWAS analyses, using region-level methylation averages as the phenotype vector. On the genotype level, we considered all genotyped SNPs among the 645 accessions that had a minor allele frequency (MAF) > 5% (1,813,837 SNPs). Using this strategy, we identified 16,083 regions in CG, 8,102 in CHG, and 19,388 in CHH context that yielded at least one SNP association with significance levels that passed the Bonferroni-corrected significance threshold at α = 0.05 (threshold = 2.8×10^−8^).

Combining these results in a meta-analysis revealed that many of these associations marked region-proximal SNP locations in *cis* for CG (Supplementary Figure 5), CHG and CHH (Figure 6) methylation contexts. Intriguingly, a few also tagged several recurring association signals in *trans* (Figure 6). For CHG DMRs, we found two recurrent associations with markers on chromosomes 3 and 5, respectively (Figure 6A,D,E). Chr3:4,496,047 was found to be associated with mean methylation rates in 97 DMRs with consistently decreased methylation in accessions carrying the alternative allele (Figure 6F,G). The SNP was located in close proximity to the *miR823A* locus, which encodes the primary miRNA (pri-miRNA) of microRNA (miRNA) 823A, known to target *CMT3. CMT3* is the main methyltransferase depositing CHG methylation (Lindroth et al., 2001) and via its chromodomain provides a direct link between DNA methylation and H3K9 dimethylation at constitutive heterochromatin in a concerted feedback loop with KRYPTONITE (KYP) and *SUPPRESSOR OF VARIATION 3-9 HOMOLOGUE PROTEIN 5/6* (*SUVH5/6*) (Du et al., 2012; Jackson et al., 2002; Du et al., 2014; Ebbs and Bender, 2006).

**Figure 6:**
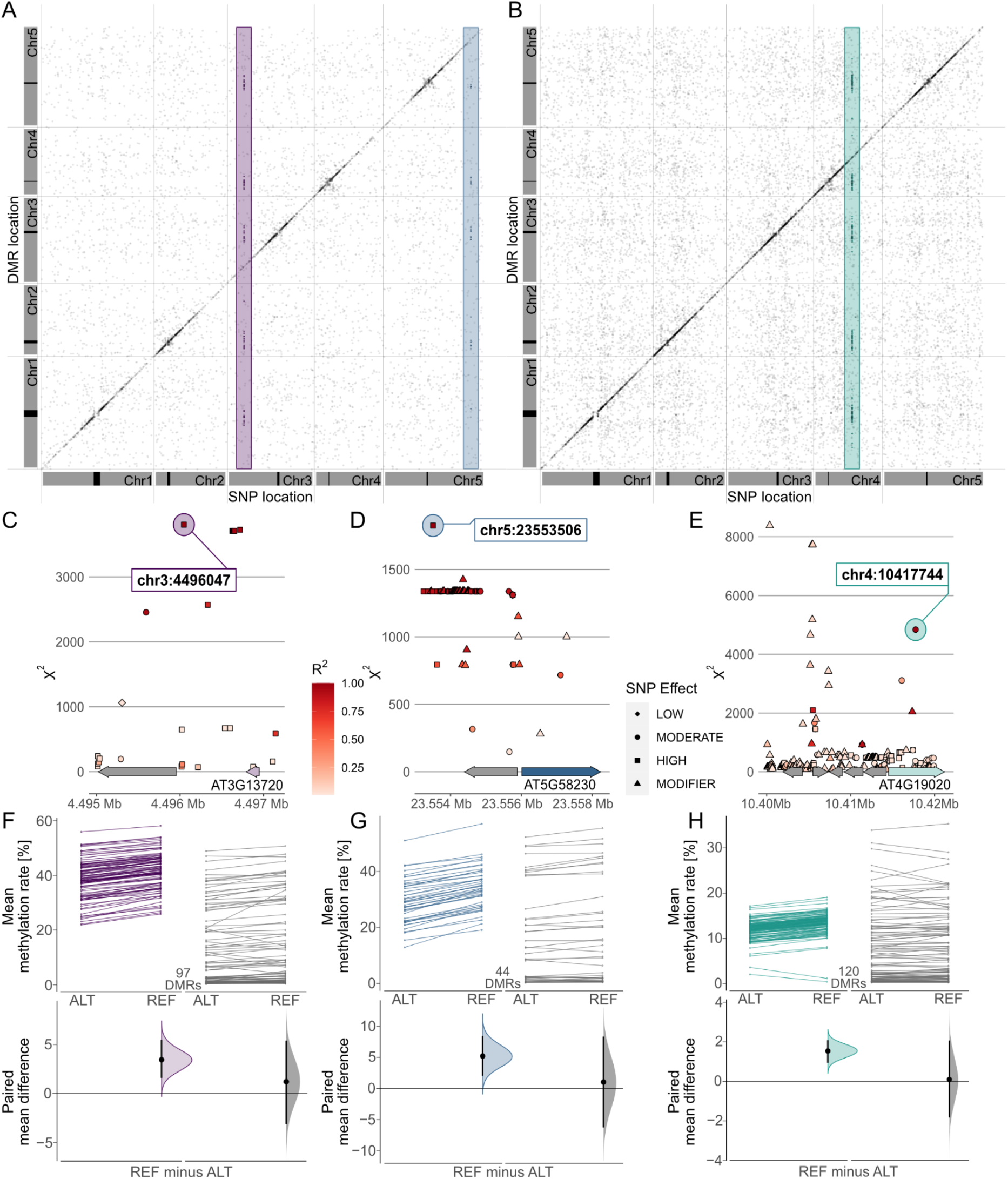
Genome Wide Association (GWA) signals recurrently emerge from differential methylation found across the *A. thaliana* 1001 Methylomes panel (Kawakatsu et al., 2016). GWA analyses on region-level methylation rate averages reveal recurrent signals in CHG (**A**) and CHH (**B**) methylation contexts. For each DMR, only top ranked SNPs that pass the Bonferroni corrected significance threshold at α = 0.05 are included, based on the number of SNP markers available across all 646 *A. thaliana* accessions used in the study (1,813,837 SNPs with minor allele frequency (MAF) > 5%, p < 2.8×10^−8^). **C-E)** Genomic loci of recurrent trans-acting SNPs highlighted in (**A**) and (**B**). **F-H**) Effect sizes of SNPs highlighted in (**A**) and (**B**) on methylation rates in regions underlying the SNP association are shown as slopegraphs and bootstrap estimates for carriers of the alternative (ALT) and reference (REF) alleles, respectively. In each plot, an equally sized set of randomly selected DMRs (in gray) is included for comparison.

The second CHG-DMR-associated locus, Chr5:23,553,506, showed association in 44 DMRs, similarly with relatively lower methylation levels in accessions carrying the alternative allele. The SNP resolved a genomic region upstream of *MULTICOPY SUPPRESSOR OF IRA1* (*MSI1*) and appeared to be in linkage with several markers in that region (Figure 6D). MSI1 acts in the evolutionarily conserved Retinoblastoma pathway and is implicated in genomic imprinting of the *FLOWERING WAGENINGEN* (*FWA*) and *FERTILIZATION INDEPENDENT SEED 2* (*FIS2*) in *A. thaliana* via direct interaction with *RETINOBLASTOMA RELATED 1* (*RBR1*) protein (Jullien et al., 2008). Interestingly, both RBR1 and MSI1 have also been identified as integral components of the *Arabidopsis* DREAM complex (Ning et al., 2020).

In the CHH context, our finding of a recurrent association at Chr5:10417744 (Figure 6B,E) confirms a recent study that identified natural alleles of *CHROMOMETHYLASE2* (*CMT2*) as determinants of CHH methylation (Sasaki et al., 2019). The association signal emerged from 120 DMRs, showing a general trend of lower methylation in accessions carrying the alternative allele, with the exception of two regions of overall low CHH methylation (Figure 6H). In summary, this shows that MethylScore provides context-specific DMRs from large WGBS datasets that can be used to determine specific genome-epigenotype associations.

## Discussion

Genome-wide studies of DNA methylation typically aim to reveal differences in DNA methylation between samples or groups of samples. This includes identification of phenotypically relevant epialleles, finding epigenetically controlled regulatory loci, or assessing naturally occurring epigenetic variation. Here, we have presented MethylScore, a pipeline for the identification and characterization of differentially methylated loci specifically from plant WGBS data. We built MethylScore in such a way as to make it accessible to a broad community of plant researchers, requiring minimal computational background; provided with alignment files or methylation metrics files and a simple sample information table, MethylScore can be run with a single command line.

Currently available DMR callers are mostly built on assumptions based on mammalian DNA methylation, including the sequence context, genomic distribution and frequency of methylated cytosines, and the average methylation rate. Plant DNA methylation violates many of these assumptions. Moreover, most DMR callers typically follow one of three general strategies, DMP clustering, genome tiling, or pre-defining regions of interest, each with their respective caveats. DMP clustering first tests for DMPs and merges spatially adjacent loci to define DMRs. Examples of such tools are *DSS* (Feng et al., 2014), *Bisulfighter* (Saito et al., 2014), *MOABS* (Sun et al., 2014), *BSmooth* (Hansen et al., 2012), and *methylpy* (Ziller et al., 2013; Schultz et al., 2015). DMP analysis typically involves statistical tests on each cytosine in the genome, racking up a heavy multiple testing burden. If no appropriate region-level testing is applied, this multiple testing problem is carried over to the DMRs, resulting in inappropriate false discovery rate (FDR) control caused by significance thresholding at single loci (Korthauer et al., 2018). In addition, many such approaches classify DMRs based on mere presence of DMPs without considering the directionality of methylation change. Multiple-testing limitations also apply to tiling approaches in which testable segments correspond to windows or sliding windows along the genome, implemented for example in *methylKit* (Akalin et al., 2012). The third category of DMR callers use predefined regions, selected based on existing knowledge, e.g. genome annotation features. These can be gene bodies, promoter regions, transposons, or predetermined CpG islands. While this reduces the multiple-testing problem, it ignores potentially relevant loci that are not included in the predefined set, yet makes tools such as *BiSeq* (Hebestreit et al., 2013) suited for targeted sequencing approaches such as reduced-representation bisulfite sequencing (RRBS) (Meissner et al., 2005).

All of the above methods have additional downsides when applied to *plant* WGBS data. Methylation occurring in all sequence contexts in plants aggravates the DMP multiple testing problem. Some approaches, including *BSmooth* (Hansen et al., 2012), *BiSeq* (Hebestreit et al., 2013), and *dmrseq* (Korthauer et al., 2018), moreover assume local correlation between spatially adjacent methylated cytosines to compute smoothed methylation estimates. Despite the presence of CHH islands that have been reported for some species (Zemach et al., 2010; Gent et al., 2013), this assumption primarily holds true for studies in mammalian genomes with high density of methylated CpG regions. It is, however, less applicable in plants and potentially affects the performance of these methods in determining genomic boundaries of methylated regions. Similarly, window- or tile-based approaches are best applicable when methylation is evenly and densely distributed. However, in *A. thaliana* and other plant genomes, the overwhelming share of methylated cytosines are concentrated in only a small fraction of the genome. Window-based approaches therefore cause a substantial penalty in statistical power when controlling for FDR, because many of the statistical tests lack biological relevance in absence of methylation in the windows that are tested.

### *MethylScore does not require* a priori *information on DNA methylation parameters or dataset structures*

In an unsupervised training step, MethylScore identifies the methylation rate distribution per sequence context in the actual data, followed by classifying the genome into states of high and low methylation based on these training parameters. Alternatively, users can decide to train MethylScore on a reference sample and apply the training parameters on all other samples in their dataset. Only those regions are tested for differential methylation in which high-methylation states differ across samples, and so our pipeline avoids testing regions with no or very little DNA methylation variation across samples, which drastically reduces the number of necessary statistical tests compared to other methods.

As shown in the test cases above, MethylScore can handle small datasets with only few samples to very large datasets with hundreds of WGBS runs. Unless users decide to provide information on replicate groups, the pipeline applies a population-scale analysis, clustering samples into groups with similar methylation rates for each testable genomic segment. This has two effects: first, MethylScore is thus able to reveal hidden sample structures that researchers might not be aware of at the start of the study. Second, substantial within-group variance, e.g. within a group of replicates of the same treatment condition, will become apparent in the DMR output and will not be masked by a forced *a priori* grouping of replicates.

### A modular pipeline that can be integrated with other WGBS analysis pipelines

MethylScore is designed as a post-alignment, secondary analysis workflow starting from either alignments in *bam* format or pre-existing single-cytosine metrics in *bedGraph* format, in contrast to existing end-to-end solutions such as *wg-blimp* (Wöste et al., 2020) or *PiGx bsseq* (Wurmus et al., 2018). However, its modular design also allows for tight integration with primary analysis pipelines such as *nf-core methylseq* (Ewels et al., 2020), *snakePipes WGBS* (Bhardwaj et al., 2019), or *EpiDiverse Toolkit* (Nunn et al., 2021). These pipelines take raw BS-sequencing reads as an input and perform extensive quality control, along with trimming and read mapping, which is highly advised to rule out or mitigate potential biases arising from experimental factors such as library preparation protocols, adapter content, and sequence duplication levels, before attempting to identify DMRs.

### Limitations and shortcomings of the pipeline

MethylScore is extensively parameterised. While we show that the default parameter set is suitable on data derived from *A. thaliana* and *O. sativa*, analysis of other species with different methylation characteristics might require a parameter sweep for optimal results.

## Outlook

Despite recent advances in detecting chemically modified bases in long-read sequencing data, WGBS remains the current standard for the analysis of whole-genome DNA methylation profiles. This applies even more to plants, for which the complexity and diversity of base modifications poses major hurdles to long-read-based technologies. Even if these technical issues will be overcome in future, statistical analysis of differential methylation to determine relevant epiallelic loci will remain a key challenge in this type of studies.

## Materials and methods

### Pipeline strategy

When provided with alignments as input, MethylScore can first merge and coordinate sort technical replicates using *samtools* v1.9 (Li et al., 2009) and subsequently deduplicates reads arisen from PCR amplification using *picard* v2.26.8 (Picard toolkit, 2019). To enable parallel processing, alignments are split by chromosome to concurrently extract single cytosine metrics using *MethylDackel* v0.4.0 (*https://github.com/dpryan79/MethylDackel*). The resulting read counts are collected for all samples and tabulated into an intermediate, custom-format table containing chromosomal position, sequence context, strand information, unmethylated and methylated read counts, as well as derived metrics such as coverage and methylation rate per cytosine. This information is collected in the genome matrix, a single file containing all of the above information for every sample and for each cytosine covered in at least one sample.

Using this genome-wide information, a two-state HMM is trained iteratively using the Baum-Welch algorithm for each of the samples as described by (Molaro et al., 2011) with two key adaptations. First, to account for the fact that, in contrast to DNA methylation in mammals, plant genomes often only harbor DNA methylation in a small fraction of the genome, the original implementation was adapted to detect *hyper*-methylated rather than *hypo*-methylated regions, effectively achieved by inverting the methylation rates. Second, we adapted the extension to a total of three sequence contexts, allowing the model to learn distinct parameters independently for CG, CHG and CHH sequence contexts for both states. This results in a total of six distributions with corresponding emission probabilities as well as transition probabilities between the methylated and unmethylated states.

Given the parameter estimates after model convergence for a maximum of 30 iterations, the most likely path is determined by Posterior Decoding and used for probabilistic segmentation of the genome into methylated regions (MRs). During this step, MR breakpoints are introduced at either chromosome boundaries or when a parameter is exceeded that defines the maximum sequence length devoid of covered cytosine positions (named “desert size”, default value 100 bp). This user-defined parameter prevents extending MRs over 100 bp without any methylation information. Having segmented the genome into MRs, MethylScore then applies two complementary population-scale approaches to reduce complexity of the statistical testing framework. First, candidate regions for significance testing are restricted to those with a user-defined change in MR frequency across samples (default: 20% change of MR frequency within 30 bp), indicative of natural region boundaries that are widespread in the sample population. Additional filters can be set to avoid testing short or unreliable MRs by requiring a minimum number of covered cytosines (default: 5) and a lower bound for coverage at those sites (default: 3). The second approach used by MethylScore to further decrease the number of statistical tests is to employ *k*-means clustering in order to group samples by mean methylation rates within any given MR. For each candidate region, cluster centers are searched to minimize the within-group variance. The value of *k* is iteratively incremented starting from *k* = 2, until the pairwise comparison of all cluster centers results in a methylation difference of less than a user-defined value, in which case the previous *k* is chosen. This parameter lets the user choose a target minimum methylation difference between groups of samples and thus can be effectively used to discard likely biologically irrelevant DMRs with only a few percentage points methylation difference.

Remaining regions that fulfill aforementioned criteria are tested for differential methylation using a beta-binomial-based test; resulting p-values are subsequently corrected for multiple tests as described in (Becker et al., 2011).

### Implementation in Nextflow

To achieve high standards of reproducibility and portability of pipeline execution across different compute infrastructures, the workflow manager Nextflow (Di Tommaso et al., 2017) was chosen to implement MethylScore. Nextflow enables containerised execution using a prebuilt Docker image (available at https://quay.io/repository/beckerlab/methylscore), that contains all necessary dependencies including perl libraries, the pre-compiled HMM implementation, and version-pinned bioconda packages. Furthermore, Nextflow ensures efficient use of available computational infrastructure by straightforward parallelisation of tasks, following a scatter-gather strategy.

### Re-analysis of published datasets

WGBS reads were retrieved from the ENA and GEO repositories PRJEB26932 (Wibowo et al., 2016), PRJEB12413 (Tedeschi et al., 2019), GSE137754 (Ning et al., 2020) GSE42410 (Stroud et al., 2013a) and GSE171414 (Zhang et al., 2021)

Raw data were trimmed and mapped to the TAIR10 (Lamesch et al., 2012) reference genome for *A. thaliana* data and the IGRSP-1.0 genome (Kawahara et al., 2013) for *O. sativa* spp. *japonica* data using v1.6 of the *nf-core methylseq* pipeline (Ewels et al., 2021). Standard settings for Illumina adapter trimming, and the default aligner *bismark* (Krueger and Andrews, 2011) were used with addition of the --`relax_mismatches` preset. The rich quality control output of the pipeline was used to assess methylation bias on both read ends and affected bases were ignored for downstream analysis with MethylScore.

### 1001 Methylomes analysis

Single-cytosine metrics obtained from the *A. thaliana* 1001 Methylomes dataset (Kawakatsu et al., 2016) were used to call DMRs in pairwise comparisons between 645 accessions using Col-0 as the point of reference. Overlapping regions found in multiple comparisons were merged using the *plyranges* (v.1.14.0 (Lee et al., 2019) *reduce_ranges()* function in *R* (v4.1.2 (R Core Team, 2021) For each sequence context, methylation rates within all identified DMRs were retrieved and used to seek associations between the average methylation rate for each accession within each DMR and the underlying genotype by conducting Genome-Wide Association Studies (GWAS) using the *Limix* framework (Lippert et al., 2014) for linear mixed models using an in-house python implementation packaged into a containerised Nextflow pipeline (https://gitlab.lrz.de/beckerlab/gwas-nf).

In brief, our GWAS pipeline retrieves the average methylation rate for each accession within a given DMR and estimates population structure from all SNPs using the *limix*.*stats*.*linear_kinship* function in *Limix*. GWAS is performed via the *limix*.*qtl*.*scan* function after subsetting the full genome SNP matrix (1001 Genomes Consortium, 2016) to include all polymorphic SNPs that have a minor allele frequency (MAF) of at least 5% across the population of 645 accessions. For meta analyses, only those SNP positions are reported for which the significance level exceeds the Bonferroni corrected threshold with respect to the total number of included SNP markers (1,813,837).

### Data visualization

Figures were generated using *R* 4.1.2 (R Core Team, 2021) and *Bioconductor* 3.14 (Gentleman et al., 2004; Huber et al., 2015). Heatmaps of methylation rates were generated using the *ComplexHeatmap* library (v2.10.0 (Gu et al., 2016). Principal components were computed using the *pcaMethods* library (v1.86.0 (Stacklies et al., 2007) and visualized with *ggplot2* (v3.3.5 (Wickham, 2009) and the *ggrepel* library (v0.9.1 (Slowikowski). SNP linkage was calculated using the *snpStats* library (v1.44.0) and genomic locations were illustrated using the *gggenes* library (v0.4.1 (Wilkins). Genome wide representations of SNP associations were drawn using *ggplot2* (v3.3.5 (Wickham, 2009) and the *ggforce* (v0.3.3 (Pedersen) library for visual annotation. Slopegraphs and Bootstrap Estimation plots were generated using the *dabestr* library (v0.3.0 (Ho et al., 2019).

## Acknowledgements

We would like to thank Niklas Schandry, Eva Knoch, and Daniela Ramos Cruz for critical reading of the manuscript and helpful suggestions. The computational results presented were obtained using the CLIP cluster at Vienna BioCenter (VBC) and the BioHPC Genomics cluster housed at the Leibniz Supercomputing Centre (LRZ).

This research was supported by the Austrian Academy of Sciences (P.H., C.B., R.P.); the European Union’s Horizon 2020 research and innovation programme via the European Research Council (ERC) Grant agreement No. 716823 “FEAR-SAP” (P.H., C.B.), DFG project ERA-CAPS AUREATE (D.W.), and via the Marie Sklodowska-Curie ETN ‘EpiDiverse’ (A.N., C.B.), grant agreement no. 764965 “Epidiverse”. F.J. acknowledges support from the Technical University of Munich-Institute for Advanced Study funded by the German Excellent Initiative and the European Seventh Framework Program under grant agreement no. 291763. F.J., and I.K. were supported by the SFB Sonderforschungsbereich924 of the Deutsche Forschungsgemeinschaft (DFG).

## Author contributions

P.H., J.H., F.J., D.W., and C.B. conceived the study and developed the pipeline; D.L. and C.B. supervised the work. J.H. wrote the MethylScore code; P.H. built the Nextflow pipeline. I.K. generated the epiRIL data. P.H. and A.N. analyzed the data. R.P. prepared methylation calls on the *A. thaliana* 1001 Genomes dataset. P.H. and C.B. wrote the manuscript.

## Competing Interests

The authors declare the following competing interests: JH is currently an employee of Computomics GmbH. S.J.S. is currently the CEO of and holds shares in Computomics GmbH.

## Data availability

The MethylScore pipeline is available on Github: https://github.com/Computomics/MethylScore. The pipeline that was used to conduct GWA studies is available on Gitlab: https://gitlab.lrz.de/beckerlab/gwas-nf.

## Supplemental Material

**Supplementary Figure 1:**
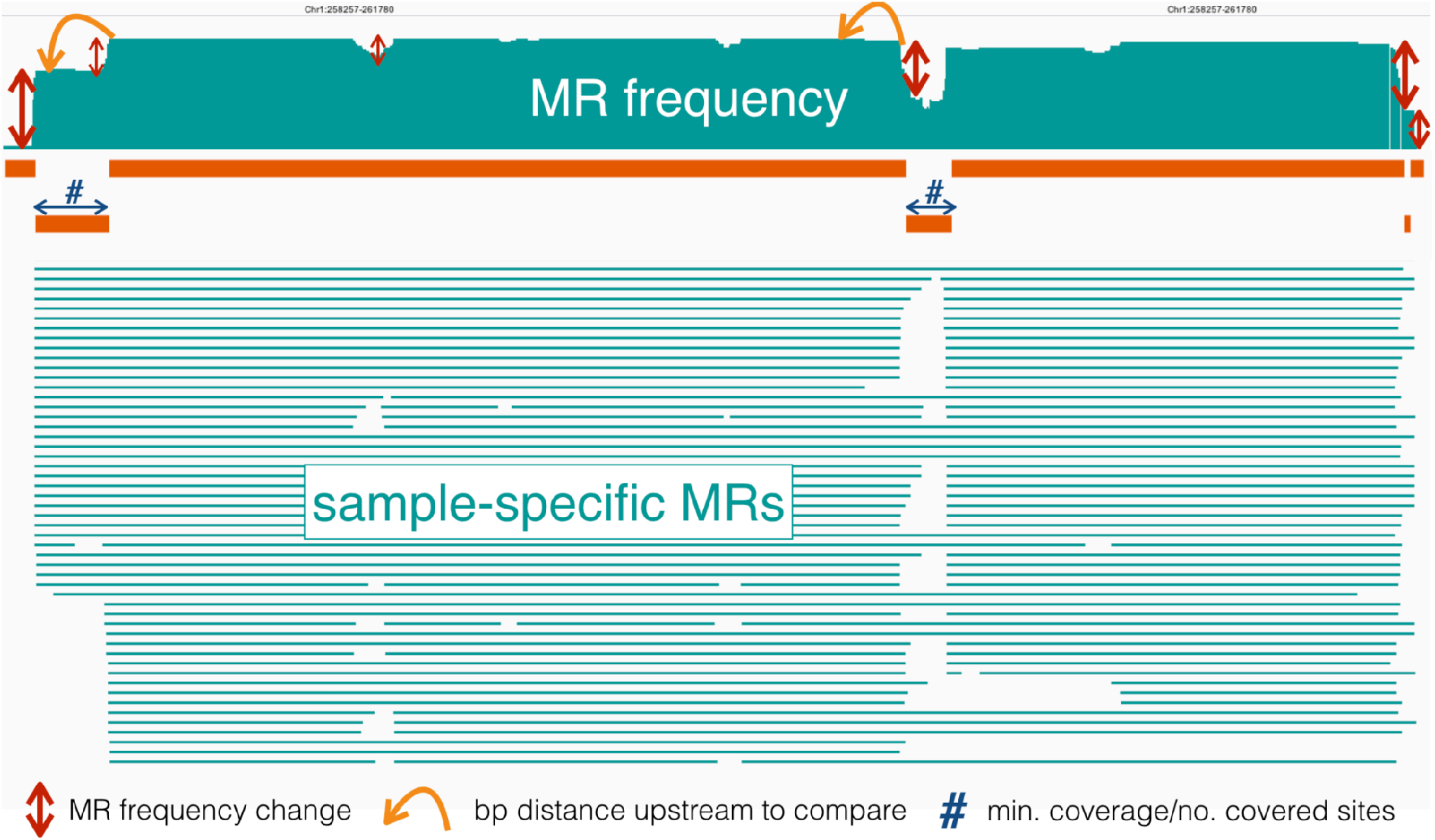
MR-based strategy to determine candidate DMRs. The top panel shows a methylation region (MR) frequency barplot along the genome, summarizing the counts of all sample-specific MRs (the MRs are displayed as horizontal petrol-colored lines in the middle). Three parameters specify the borders of candidate differentially methylated regions (DMRs; orange horizontal bars): 1) the degree of MR frequency changes along the genome (marked by red vertical arrows), 2) the upstream distance to which MR frequency is compared to (marked by orange-colored bent arrows), and 3) a minimum number and coverage of cytosines (blue arrows indicating the length of candidate DMRs).

**Supplementary Figure 2:**
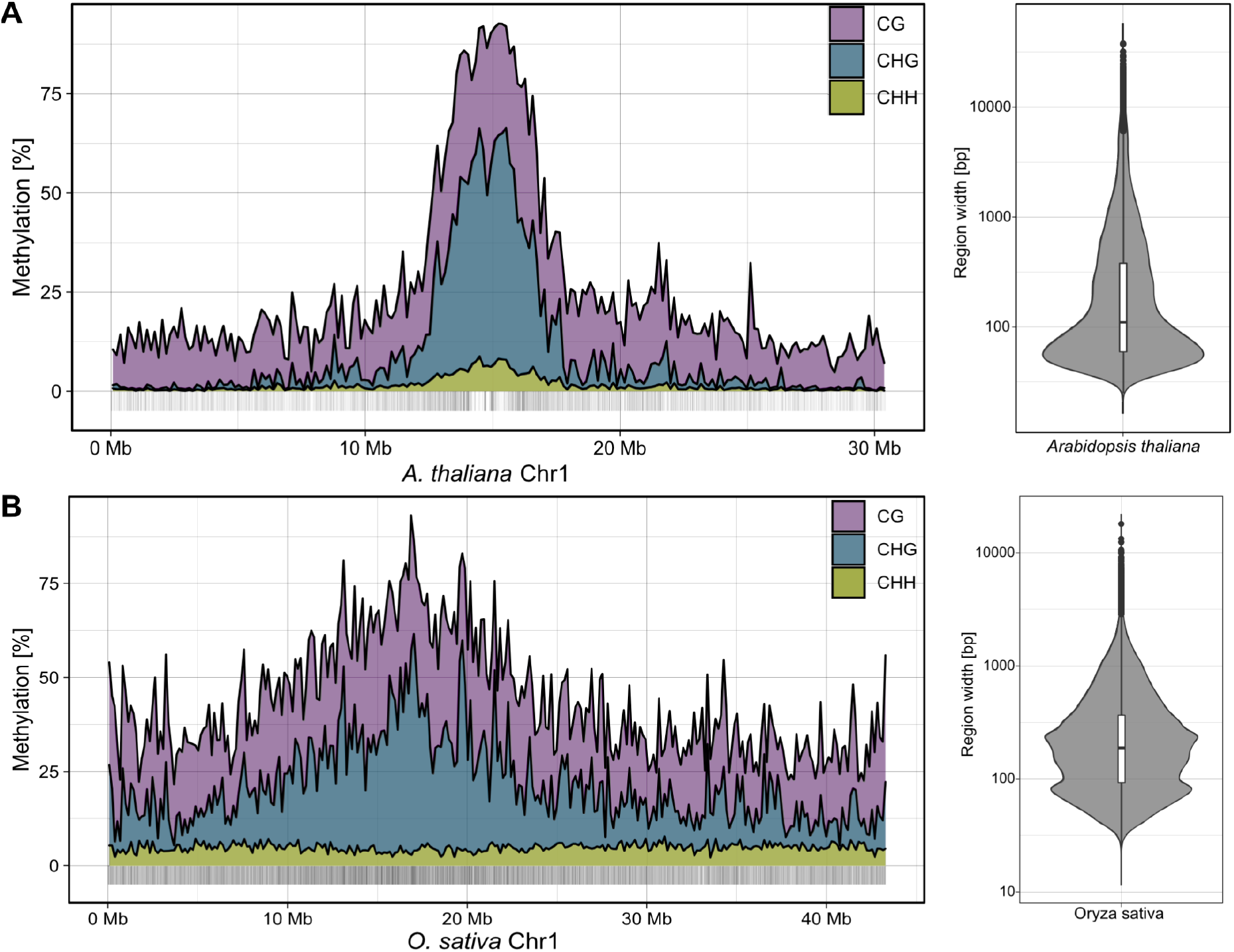
Genomic distribution of DNA methylation and methylated regions (MRs). Lineplots (left) show the distribution of average DNA methylation in 150 kb bins along single chromosomes of *A. thaliana* (**A;** data from (Kawakatsu et al., 2016) and *O. sativa* (**B;** data from (Stroud et al., 2013a). MRs segmented by MethylScore are indicated as vertical bars along the *x*-axis, with corresponding length distributions shown as violin plots (right).

**Supplementary Figure 3:**
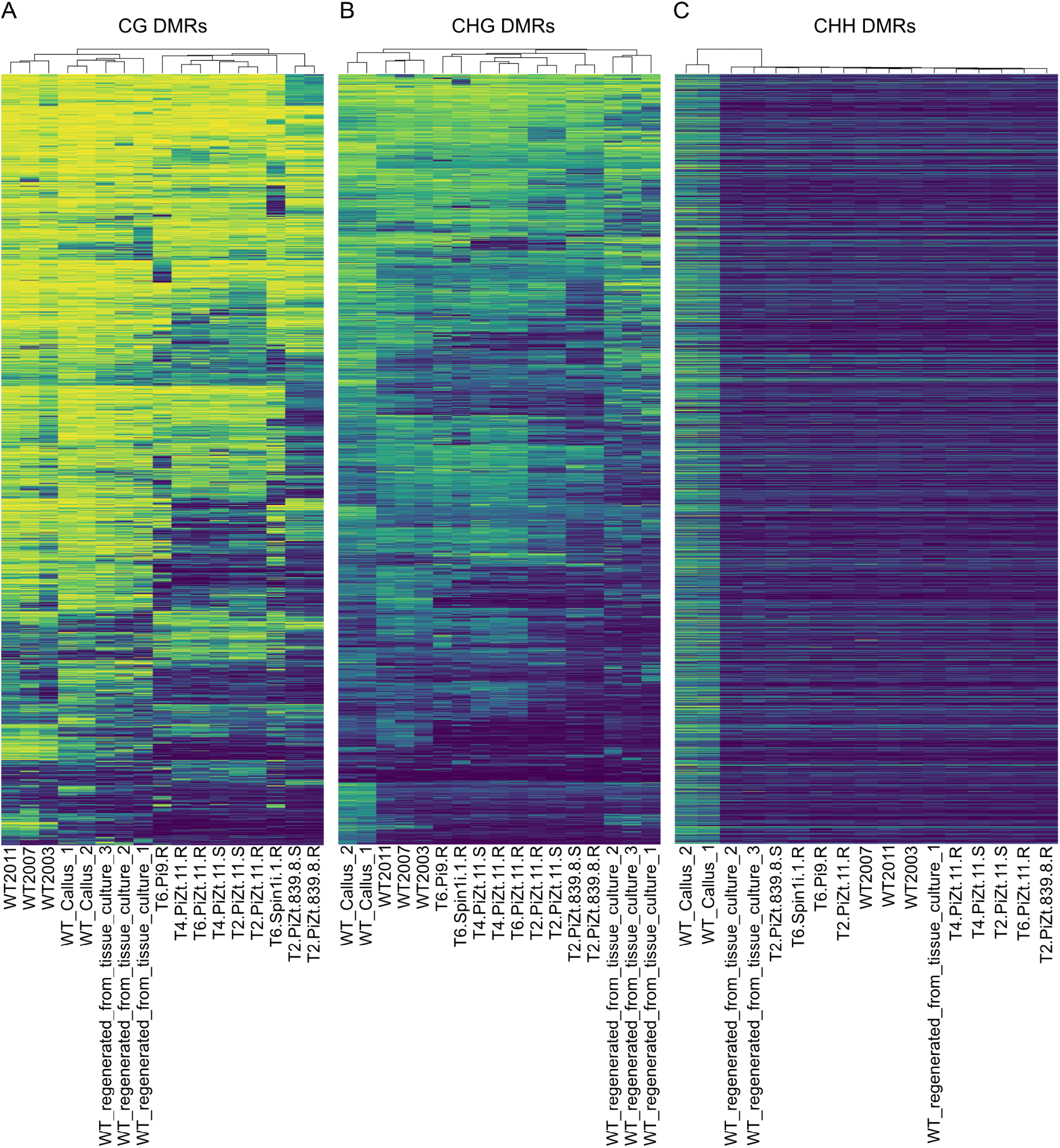
Differential methylation in rice regenerants. Heatmaps show methylation rate averages in regions identified as differentially methylated in CG, CHG, or CHH methylation contexts (from left to right) in WGBS data from different rice regenerants, callus tissue, and non-regenerated controls. Original data from (Stroud et al., 2013a), GEOaccession GSE42410.

**Supplementary Figure 4:**
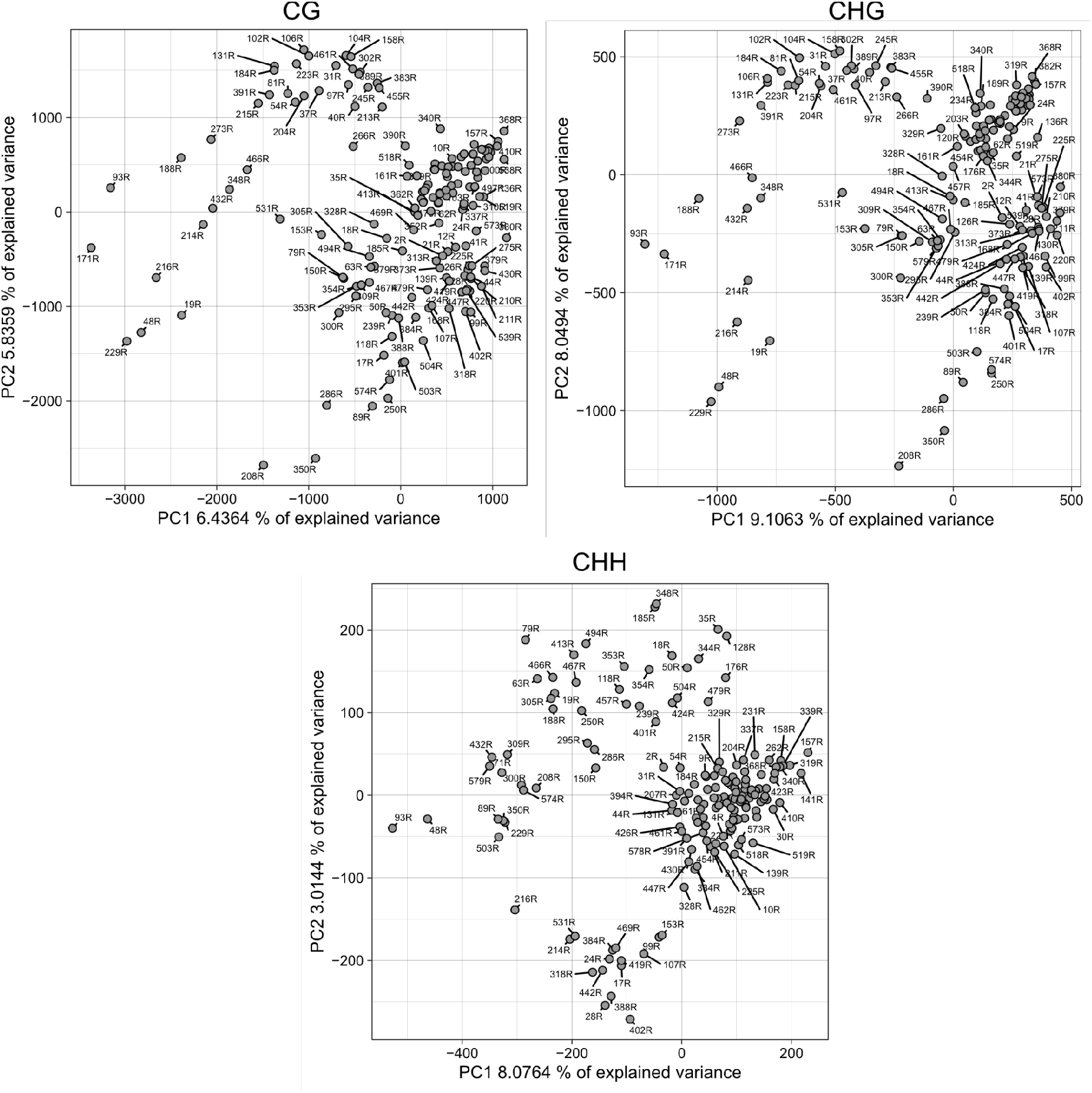
DMR identification across epigenetic recombinant inbred lines (epiRILs). Principal component analysis (PCA) of mean methylation rates within 46,323 CG-, 8,821 CHG- and 4,121 CHH-DMRs identified by MethylScore across a population of 169 epiRILs.

**Supplementary Figure 5:**
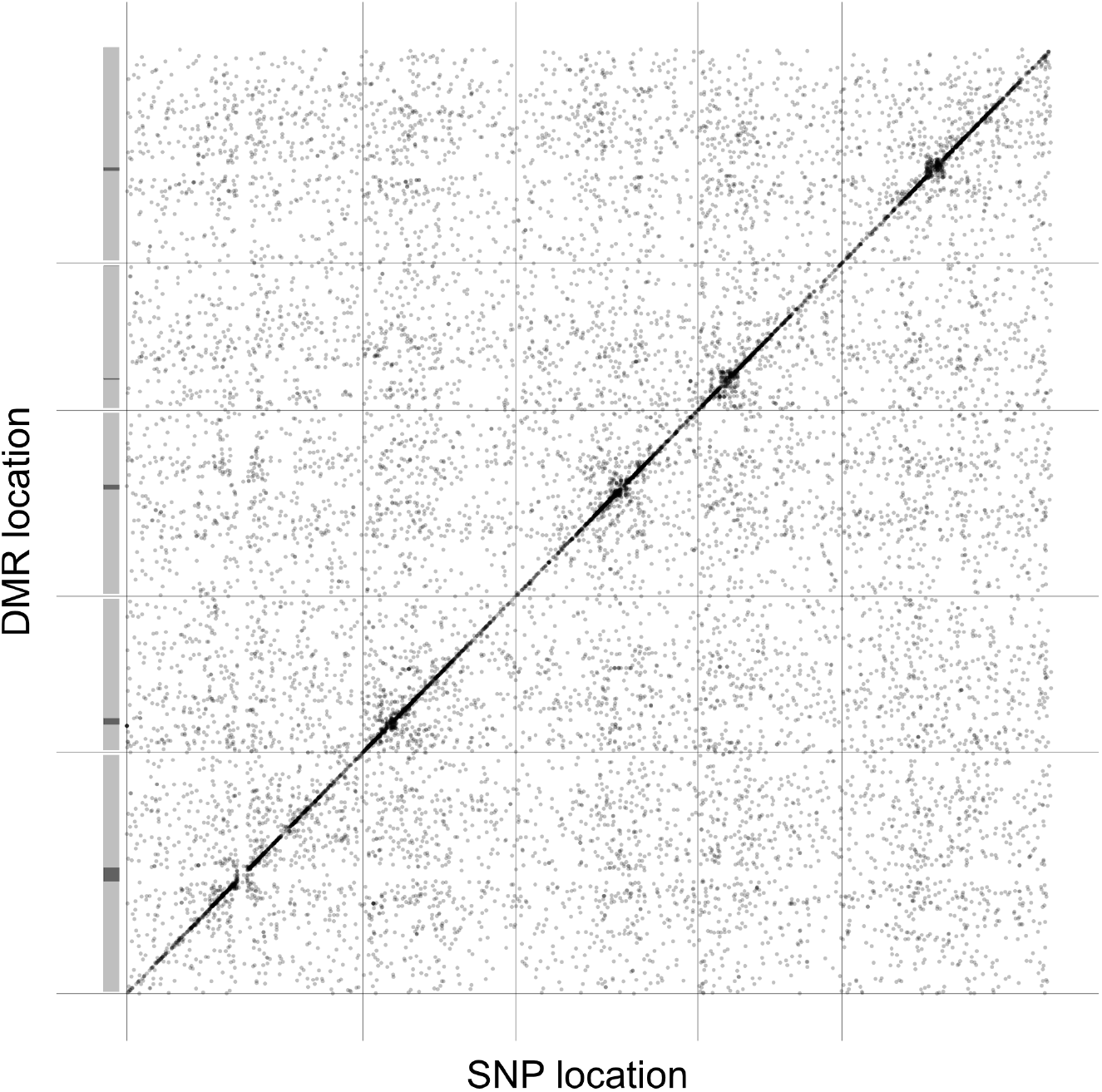
GWA analysis on region-level methylation rate averages in CG.

## References

1001 Genomes Consortium (2016). 1,135 genomes reveal the global pattern of polymorphism in Arabidopsis thaliana. Cell 166: 481–491.

Akalin, A., Kormaksson, M., Li, S., Garrett-Bakelman, F.E., Figueroa, M.E., Melnick, A., and Mason, C.E. (2012). methylKit: a comprehensive R package for the analysis of genome-wide DNA methylation profiles. Genome Biol. 13: R87.

Aufsatz, W., Florian Mette, M., van der Winden, J., Matzke, A.J.M., and Matzke, M. (2002). RNA-directed DNA methylation in Arabidopsis. Proc. Natl. Acad. Sci. U. S. A. 99: 16499–16506.

Becker, C., Hagmann, J., Müller, J., Koenig, D., Stegle, O., Borgwardt, K., and Weigel, D. (2011). Spontaneous epigenetic variation in the Arabidopsis thaliana methylome. Nature 480: 245–249.

Bhardwaj, V., Heyne, S., Sikora, K., Rabbani, L., Rauer, M., Kilpert, F., Richter, A.S., Ryan, D.P., and Manke, T. (2019). snakePipes: facilitating flexible, scalable and integrative epigenomic analysis. Bioinformatics 35: 4757–4759.

Cao, X., Aufsatz, W., Zilberman, D., Mette, M.F., Huang, M.S., Matzke, M., and Jacobsen, S.E. (2003). Role of the DRM and CMT3 methyltransferases in RNA-directed DNA methylation. Curr. Biol. 13: 2212–2217.

Cao, X. and Jacobsen, S.E. (2002). Role of the arabidopsis DRM methyltransferases in de novo DNA methylation and gene silencing. Curr. Biol. 12: 1138–1144.

Cervera, M.T., Ruiz-García, L., and Martínez-Zapater, J.M. (2002). Analysis of DNA methylation in Arabidopsis thaliana based on methylation-sensitive AFLP markers. Mol. Genet. Genomics 268: 543–552.

Cokus, S.J., Feng, S., Zhang, X., Chen, Z., Merriman, B., Haudenschild, C.D., Pradhan, S., Nelson, S.F., Pellegrini, M., and Jacobsen, S.E. (2008). Shotgun bisulphite sequencing of the Arabidopsis genome reveals DNA methylation patterning. Nature 452: 215–219.

Colomé-Tatché, M. et al. (2012). Features of the Arabidopsis recombination landscape resulting from the combined loss of sequence variation and DNA methylation. Proc. Natl. Acad. Sci. U. S. A. 109: 16240–16245.

Cubas, P., Vincent, C., and Coen, E. (1999). An epigenetic mutation responsible for natural variation in floral symmetry. Nature 401: 157–161.

Deaton, A.M. and Bird, A. (2011). CpG islands and the regulation of transcription. Genes Dev. 25: 1010–1022.

Di Tommaso, P., Chatzou, M., Floden, E.W., Barja, P.P., Palumbo, E., and Notredame, C. (2017). Nextflow enables reproducible computational workflows. Nat. Biotechnol. 35: 316–319.

Dubin, M.J. et al. (2015). DNA methylation in Arabidopsis has a genetic basis and shows evidence of local adaptation. Elife 4: e05255.

Du, J. et al. (2012). Dual binding of chromomethylase domains to H3K9me2-containing nucleosomes directs DNA methylation in plants. Cell 151: 167–180.

Du, J., Johnson, L.M., Groth, M., Feng, S., Hale, C.J., Li, S., Vashisht, A.A., Wohlschlegel, J.A., Patel, D.J., and Jacobsen, S.E. (2014). Mechanism of DNA methylation-directed histone methylation by KRYPTONITE. Mol. Cell 55: 495–504.

Ebbs, M.L. and Bender, J. (2006). Locus-specific control of DNA methylation by the Arabidopsis SUVH5 histone methyltransferase. Plant Cell 18: 1166–1176.

Eichten, S.R. et al. (2011). Heritable epigenetic variation among maize inbreds. PLoS Genet. 7: e1002372.

Ewels, P. et al. (2021). nf-core/methylseq: nf-core/methylseq version 1.6.1 [Nauseous Serpent].

Ewels, P.A., Peltzer, A., Fillinger, S., Patel, H., Alneberg, J., Wilm, A., Garcia, M.U., Di Tommaso, P., and Nahnsen, S. (2020). The nf-core framework for community-curated bioinformatics pipelines. Nat. Biotechnol. 38: 276–278.

Feng, H., Conneely, K.N., and Wu, H. (2014). A Bayesian hierarchical model to detect differentially methylated loci from single nucleotide resolution sequencing data. Nucleic Acids Res. 42: e69.

Fick, S.E. and Hijmans, R.J. (2017). WorldClim 2: new 1-km spatial resolution climate surfaces for global land areas. Int. J. Climatol. 37: 4302–4315.

Finnegan, E.J., Peacock, W.J., and Dennis, E.S. (1996). Reduced DNA methylation in Arabidopsis thaliana results in abnormal plant development. Proc. Natl. Acad. Sci. U. S. A. 93: 8449–8454.

Frommer, M., McDonald, L.E., Millar, D.S., Collis, C.M., Watt, F., Grigg, G.W., Molloy, P.L., and Paul, C.L. (1992). A genomic sequencing protocol that yields a positive display of 5-methylcytosine residues in individual DNA strands. Proc. Natl. Acad. Sci. U. S. A. 89: 1827–1831.

Gehring, M., Huh, J.H., Hsieh, T.-F., Penterman, J., Choi, Y., Harada, J.J., Goldberg, R.B., and Fischer, R.L. (2006). DEMETER DNA glycosylase establishes MEDEA polycomb gene self-imprinting by allele-specific demethylation. Cell 124: 495–506.

Gent, J.I., Ellis, N.A., Guo, L., Harkess, A.E., Yao, Y., Zhang, X., and Dawe, R.K. (2013). CHH islands: de novo DNA methylation in near-gene chromatin regulation in maize. Genome Res. 23: 628–637.

Gentleman, R.C. et al. (2004). Bioconductor: open software development for computational biology and bioinformatics. Genome Biol. 5: R80.

van der Graaf, A., Wardenaar, R., Neumann, D.A., Taudt, A., Shaw, R.G., Jansen, R.C., Schmitz, R.J., Colomé-Tatché, M., and Johannes, F. (2015). Rate, spectrum, and evolutionary dynamics of spontaneous epimutations. Proc. Natl. Acad. Sci. U. S. A. 112: 6676–6681.

Gu, Z., Eils, R., and Schlesner, M. (2016). Complex heatmaps reveal patterns and correlations in multidimensional genomic data. Bioinformatics 32: 2847–2849.

Hagmann, J., Becker, C., Müller, J., Stegle, O., Meyer, R.C., Wang, G., Schneeberger, K., Fitz, J., Altmann, T., Bergelson, J., Borgwardt, K., and Weigel, D. (2015). Century-scale methylome stability in a recently diverged Arabidopsis thaliana lineage. PLoS Genet. 11: e1004920.

Hansen, K.D., Langmead, B., and Irizarry, R.A. (2012). BSmooth: from whole genome bisulfite sequencing reads to differentially methylated regions. Genome Biol. 13: 1–10.

Hebestreit, K., Dugas, M., and Klein, H.-U. (2013). Detection of significantly differentially methylated regions in targeted bisulfite sequencing data. Bioinformatics 29: 1647–1653.

Henderson, I.R. and Jacobsen, S.E. (2007). Epigenetic inheritance in plants. Nature 447: 418–424.

Heyn, H., Moran, S., Hernando-Herraez, I., Sayols, S., Gomez, A., Sandoval, J., Monk, D., Hata, K., Marques-Bonet, T., Wang, L., and Esteller, M. (2013). DNA methylation contributes to natural human variation. Genome Res. 23: 1363–1372.

Ho, J., Tumkaya, T., Aryal, S., Choi, H., and Claridge-Chang, A. (2019). Moving beyond P values: data analysis with estimation graphics. Nat. Methods 16: 565–566.

Huber, W. et al. (2015). Orchestrating high-throughput genomic analysis with Bioconductor. Nat. Methods 12: 115–121.

Huff, J.T. and Zilberman, D. (2014). Dnmt1-independent CG methylation contributes to nucleosome positioning in diverse eukaryotes. Cell 156: 1286–1297.

Jackson, J.P., Lindroth, A.M., Cao, X., and Jacobsen, S.E. (2002). Control of CpNpG DNA methylation by the KRYPTONITE histone H3 methyltransferase. Nature 416: 556– 560.

Jaenisch, R. and Bird, A. (2003). Epigenetic regulation of gene expression: how the genome integrates intrinsic and environmental signals. Nat. Genet. 33: 245–254.

Jeltsch, A. (2006). On the enzymatic properties of Dnmt1: specificity, processivity, mechanism of linear diffusion and allosteric regulation of the enzyme. Epigenetics 1: 63– 66.

Johannes, F. et al. (2009). Assessing the impact of transgenerational epigenetic variation on complex traits. PLoS Genet. 5: e1000530.

Jühling, F., Kretzmer, H., Bernhart, S.H., Otto, C., Stadler, P.F., and Hoffmann, S. (2016). metilene: fast and sensitive calling of differentially methylated regions from bisulfite sequencing data. Genome Res. 26: 256–262.

Jullien, P.E., Mosquna, A., Ingouff, M., Sakata, T., Ohad, N., and Berger, F. (2008). Retinoblastoma and its binding partner MSI1 control imprinting in Arabidopsis. PLoS Biol. 6: e194.

Kankel, M.W., Ramsey, D.E., Stokes, T.L., Flowers, S.K., Haag, J.R., Jeddeloh, J.A., Riddle, N.C., Verbsky, M.L., and Richards, E.J. (2003). Arabidopsis MET1 cytosine methyltransferase mutants. Genetics 163: 1109–1122.

Kawahara, Y. et al. (2013). Improvement of the Oryza sativa Nipponbare reference genome using next generation sequence and optical map data. Rice 6: 4.

Kawakatsu, T. et al. (2016). Epigenomic diversity in a global collection of Arabidopsis thaliana accessions. Cell 166: 492–505.

Korthauer, K., Chakraborty, S., Benjamini, Y., and Irizarry, R.A. (2018). Detection and accurate false discovery rate control of differentially methylated regions from whole genome bisulfite sequencing. Biostatistics 20: 367–383.

Krueger, F. and Andrews, S.R. (2011). Bismark: a flexible aligner and methylation caller for Bisulfite-Seq applications. Bioinformatics 27: 1571–1572.

Lamesch, P. et al. (2012). The Arabidopsis Information Resource (TAIR): improved gene annotation and new tools. Nucleic Acids Res. 40: D1202–10.

Law, J.A. and Jacobsen, S.E. (2010). Establishing, maintaining and modifying DNA methylation patterns in plants and animals. Nat. Rev. Genet. 11: 204–220.

Lee, S., Cook, D., and Lawrence, M. (2019). plyranges: a grammar of genomic data transformation. Genome Biol. 20: 4.

Liégard, B., Baillet, V., Etcheverry, M., Joseph, E., Lariagon, C., Lemoine, J., Evrard, A., Colot, V., Gravot, A., Manzanares-Dauleux, M.J., and Jubault, M. (2019). Quantitative resistance to clubroot infection mediated by transgenerational epigenetic variation in Arabidopsis. New Phytol. 222: 468–479.

Li, H., Handsaker, B., Wysoker, A., Fennell, T., Ruan, J., Homer, N., Marth, G., Abecasis, G., Durbin, R., and 1000 Genome Project Data Processing Subgroup (2009). The Sequence Alignment/Map format and SAMtools. Bioinformatics 25: 2078– 2079.

Lindroth, A.M., Cao, X., Jackson, J.P., Zilberman, D., McCallum, C.M., Henikoff, S., and Jacobsen, S.E. (2001). Requirement of CHROMOMETHYLASE3 for maintenance of CpXpG methylation. Science 292: 2077–2080.

Lippert, C., Casale, F.P., Rakitsch, B., and Stegle, O. (2014). LIMIX: genetic analysis of multiple traits. bioRxiv: 003905.

Lippman, Z. et al. (2004). Role of transposable elements in heterochromatin and epigenetic control. Nature 430: 471–476.

Lister, R., O’Malley, R.C., Tonti-Filippini, J., Gregory, B.D., Berry, C.C., Harvey Millar, A., and Ecker, J.R. (2008). Highly integrated single-base resolution maps of the epigenome in Arabidopsis. Cell 133: 523–536.

Liu, G., Xia, Y., Liu, T., Dai, S., and Hou, X. (2018). The DNA methylome and association of differentially methylated regions with differential gene expression during heat stress in Brassica rapa. Int. J. Mol. Sci. 19.

MacQueen, J. (1967). Some methods for classification and analysis of multivariate observations. In Proceedings of the Fifth Berkeley Symposium on Mathematical Statistics and Probability, Volume 1: Statistics (University of California Press: Berkeley, CA), pp. 281–298.

Manning, K., Tör, M., Poole, M., Hong, Y., Thompson, A.J., King, G.J., Giovannoni, J.J., and Seymour, G.B. (2006). A naturally occurring epigenetic mutation in a gene encoding an SBP-box transcription factor inhibits tomato fruit ripening. Nat. Genet. 38: 948–952.

Manolio, T.A. et al. (2009). Finding the missing heritability of complex diseases. Nature 461: 747–753.

Matzke, M.A. and Mosher, R.A. (2014). RNA-directed DNA methylation: an epigenetic pathway of increasing complexity. Nat. Rev. Genet. 15: 394–408.

Meissner, A., Gnirke, A., Bell, G.W., Ramsahoye, B., Lander, E.S., and Jaenisch, R. (2005). Reduced representation bisulfite sequencing for comparative high-resolution DNA methylation analysis. Nucleic Acids Res. 33: 5868–5877.

Miura, A., Yonebayashi, S., Watanabe, K., Toyama, T., Shimada, H., and Kakutani, T. (2001). Mobilization of transposons by a mutation abolishing full DNA methylation in Arabidopsis. Nature 411: 212–214.

Miura, K., Agetsuma, M., Kitano, H., Yoshimura, A., Matsuoka, M., Jacobsen, S.E., and Ashikari, M. (2009). A metastable DWARF1 epigenetic mutant affecting plant stature in rice. Proc. Natl. Acad. Sci. U. S. A. 106: 11218–11223.

Molaro, A., Hodges, E., Fang, F., Song, Q., McCombie, W.R., Hannon, G.J., and Smith, A.D. (2011). Sperm methylation profiles reveal features of epigenetic inheritance and evolution in primates. Cell 146: 1029–1041.

Niederhuth, C.E. et al. (2016). Widespread natural variation of DNA methylation within angiosperms. Genome Biol. 17: 194.

Ning, Y.-Q., Liu, N., Lan, K.-K., Su, Y.-N., Li, L., Chen, S., and He, X.-J. (2020). DREAM complex suppresses DNA methylation maintenance genes and precludes DNA hypermethylation. Nat Plants 6: 942–956.

Nunn, A., Can, S.N., Otto, C., Fasold, M., Díez Rodríguez, B., Fernández-Pozo, N., Rensing, S.A., Stadler, P.F., and Langenberger, D. (2021). EpiDiverse Toolkit: a pipeline suite for the analysis of bisulfite sequencing data in ecological plant epigenetics. NAR Genom Bioinform 3: qab106.

Ong-Abdullah, M. et al. (2015). Loss of Karma transposon methylation underlies the mantled somaclonal variant of oil palm. Nature 525: 533–537.

Papareddy, R.K., Páldi, K., Smolka, A.D., Hüther, P., Becker, C., and Nodine, M.D. (2021). Repression of CHROMOMETHYLASE 3 prevents epigenetic collateral damage in Arabidopsis. eLife 10: e69396.

Pedersen, T.L. ggforce: Accelerating ggplot2 (Github).

Picard toolkit (2019). Broad Institute, GitHub repository.

Pignatta, D., Novitzky, K., Satyaki, P.R.V., and Gehring, M. (2018). A variably imprinted epiallele impacts seed development. PLoS Genet. 14: e1007469.

Quadrana, L., Etcheverry, M., Gilly, A., Caillieux, E., Madoui, M.-A., Guy, J., Bortolini Silveira, A., Engelen, S., Baillet, V., Wincker, P., Aury, J.-M., and Colot, V. (2019). Transposition favors the generation of large effect mutations that may facilitate rapid adaption. Nat. Commun. 10: 3421.

R Core Team (2021). R: A Language and Environment for Statistical Computing.

Riddle, N.C. and Richards, E.J. (2002). The control of natural variation in cytosine methylation in Arabidopsis. Genetics 162: 355–363.

Ronemus, M.J., Galbiati, M., Ticknor, C., Chen, J., and Dellaporta, S.L. (1996). Demethylation-induced developmental pleiotropy in Arabidopsis. Science 273: 654–657.

Saito, Y., Tsuji, J., and Mituyama, T. (2014). Bisulfighter: accurate detection of methylated cytosines and differentially methylated regions. Nucleic Acids Res. 42: e45.

Sasaki, E., Kawakatsu, T., Ecker, J.R., and Nordborg, M. (2019). Common alleles of CMT2 and NRPE1 are major determinants of CHH methylation variation in Arabidopsis thaliana. PLoS Genet. 15: e1008492.

Saze, H., Mittelsten Scheid, O., and Paszkowski, J. (2003). Maintenance of CpG methylation is essential for epigenetic inheritance during plant gametogenesis. Nat. Genet. 34: 65–69.

Schmitz, R.J., Schultz, M.D., Lewsey, M.G., O’Malley, R.C., Urich, M.A., Libiger, O., Schork, N.J., and Ecker, J.R. (2011). Transgenerational epigenetic instability is a source of novel methylation variants. Science 334: 369–373.

Schmitz, R.J., Schultz, M.D., Urich, M.A., Nery, J.R., Pelizzola, M., Libiger, O., Alix, A., McCosh, R.B., Chen, H., Schork, N.J., and Ecker, J.R. (2013). Patterns of population epigenomic diversity. Nature 495: 193–198.

Schultz, M.D. et al. (2015). Human body epigenome maps reveal noncanonical DNA methylation variation. Nature 523: 212–216.

Shen, X., De Jonge, J., Forsberg, S.K.G., Pettersson, M.E., Sheng, Z., Hennig, L., and Carlborg, Ö. (2014). Natural CMT2 variation is associated with genome-wide methylation changes and temperature seasonality. PLoS Genet. 10: e1004842.

Slowikowski, K. ggrepel: Repel overlapping text labels away from each other (Github).

Srivastava, A., Karpievitch, Y.V., Eichten, S.R., Borevitz, J.O., and Lister, R. (2019). HOME: a histogram based machine learning approach for effective identification of differentially methylated regions. BMC Bioinformatics 20: 253.

Stacklies, W., Redestig, H., Scholz, M., Walther, D., and Selbig, J. (2007). pcaMethods -a bioconductor package providing PCA methods for incomplete data. Bioinformatics 23: 1164–1167.

Storey, J.D. and Tibshirani, R. (2003). Statistical significance for genomewide studies. Proc. Natl. Acad. Sci. U. S. A. 100: 9440–9445.

Stroud, H., Ding, B., Simon, S.A., Feng, S., Bellizzi, M., Pellegrini, M., Wang, G.-L., Meyers, B.C., and Jacobsen, S.E. (2013a). Plants regenerated from tissue culture contain stable epigenome changes in rice. eLife 2: e00354.

Stroud, H., Do, T., Du, J., Zhong, X., Feng, S., Johnson, L., Patel, D.J., and Jacobsen, S.E. (2014). Non-CG methylation patterns shape the epigenetic landscape in Arabidopsis. Nat. Struct. Mol. Biol. 21: 64–72.

Stroud, H., Greenberg, M.V.C., Feng, S., Bernatavichute, Y.V., and Jacobsen, S.E. (2013b). Comprehensive analysis of silencing mutants reveals complex regulation of the Arabidopsis methylome. Cell 152: 352–364.

Sun, D., Xi, Y., Rodriguez, B., Park, H.J., Tong, P., Meong, M., Goodell, M.A., and Li, W. (2014). MOABS: model based analysis of bisulfite sequencing data. Genome Biol. 15: R38.

Tedeschi, F., Rizzo, P., Huong, B.T.M., Czihal, A., Rutten, T., Altschmied, L., Scharfenberg, S., Grosse, I., Becker, C., Weigel, D., Bäumlein, H., and Kuhlmann, M. (2019). EFFECTOR OF TRANSCRIPTION factors are novel plant-specific regulators associated with genomic DNA methylation in Arabidopsis. New Phytol. 221: 261–278.

Vaughn, M.W. et al. (2007). Epigenetic natural variation in Arabidopsis thaliana. PLoS Biol. 5: e174.

Wassenegger, M., Heimes, S., and Sänger, H.L. (1994). An infectious viroid RNA replicon evolved from an in vitro-generated non-infectious viroid deletion mutant via a complementary deletion in vivo. EMBO J. 13: 6172–6177.

Wibowo, A. et al. (2018). Partial maintenance of organ-specific epigenetic marks during plant asexual reproduction leads to heritable phenotypic variation. Proc. Natl. Acad. Sci. U. S. A. 115: E9145–E9152.

Wibowo, A., Becker, C., Marconi, G., Durr, J., Price, J., Hagmann, J., Papareddy, R., Putra, H., Kageyama, J., Becker, J., Weigel, D., and Gutierrez-Marcos, J. (2016). Hyperosmotic stress memory in Arabidopsis is mediated by distinct epigenetically labile sites in the genome and is restricted in the male germline by DNA glycosylase activity. eLife 5: e13546.

Wickham, H. (2009). ggplot2: Elegant Graphics for Data Analysis (Springer, New York, NY). Wilkins, D. gggenes: Draw gene arrow maps in ggplot2 (Github).

Wöste, M., Leitão, E., Laurentino, S., Horsthemke, B., Rahmann, S., and Schröder, C. (2020). wg-blimp: an end-to-end analysis pipeline for whole genome bisulfite sequencing data. BMC Bioinformatics 21: 169.

Wurmus, R., Uyar, B., Osberg, B., Franke, V., Gosdschan, A., Wreczycka, K., Ronen, J., and Akalin, A. (2018). PiGx: reproducible genomics analysis pipelines with GNU Guix. Gigascience 7.

Zemach, A., McDaniel, I.E., Silva, P., and Zilberman, D. (2010). Genome-wide evolutionary analysis of eukaryotic DNA methylation. Science 328: 916–919.

Zemach, A., Yvonne Kim, M., Hsieh, P.-H., Coleman-Derr, D., Eshed-Williams, L., Thao, K., Harmer, S.L., and Zilberman, D. (2013). The Arabidopsis nucleosome remodeler DDM1 allows DNA methyltransferases to access H1-containing heterochromatin. Cell 153: 193–205.

Zhang, Y., Jang, H., Xiao, R., Kakoulidou, I., Piecyk, R.S., Johannes, F., and Schmitz, R.J. (2021). Heterochromatin is a quantitative trait associated with spontaneous epiallele formation. Nat. Commun. 12: 1–13.

Zhong, S., Fei, Z., Chen, Y.-R., Zheng, Y., Huang, M., Vrebalov, J., McQuinn, R., Gapper, N., Liu, B., Xiang, J., Shao, Y., and Giovannoni, J.J. (2013). Single-base resolution methylomes of tomato fruit development reveal epigenome modifications associated with ripening. Nat. Biotechnol. 31: 154–159.

Ziller, M.J., Gu, H., Müller, F., Donaghey, J., Tsai, L.T.-Y., Kohlbacher, O., De Jager, P.L., Rosen, E.D., Bennett, D.A., Bernstein, B.E., Gnirke, A., and Meissner, A. (2013). Charting a dynamic DNA methylation landscape of the human genome. Nature 500: 477–481.

